# Keratinocyte-tethering modification for biologics enables location-precise treatment in mouse vitiligo

**DOI:** 10.1101/2022.02.28.482387

**Authors:** Ying-Chao Hsueh, Yuzhen Wang, Rebecca L. Riding, Donna E. Catalano, Yu-Jung Lu, Jillian M. Richmond, Don L. Siegel, Mary Rusckowski, John R. Stanley, John E. Harris

## Abstract

Despite the central role of IFNγ in vitiligo pathogenesis, systemic IFNγ neutralization is an impractical treatment option due to strong immunosuppression. However, most vitiligo patients present with less than 20% affected body surface area, which provides an opportunity for localized treatments that avoid systemic side effects. After identifying keratinocytes as key cells that amplify IFNγ signaling during vitiligo, we hypothesized that tethering an IFNγ neutralizing antibody to keratinocytes would limit anti-IFNγ effects to the treated skin for the localized treatment. To that end, we developed a bispecific antibody (BsAb) capable of blocking IFNγ signaling while binding to desmoglein expressed by keratinocytes. We characterized the effect of the BsAb *in vitro*, *ex vivo*, and in a mouse model of vitiligo. SPECT/CT biodistribution and serum assays after local footpad injection revealed that the BsAb had improved skin retention, faster elimination from the blood, and less systemic IFNγ inhibition than the non-tethered version. Furthermore, the BsAb conferred localized protection almost exclusively to the treated footpad during vitiligo that was not possible by local injection of the non-tethered anti-IFNγ antibody. Thus, keratinocyte-tethering proved effective while significantly diminishing off-tissue effects of IFNγ blockade, offering a new treatment strategy for localized skin diseases, including vitiligo.

## INTRODUCTION

Vitiligo is characterized by progressive demarcated white spots devoid of melanocytes in the lesional skin, caused by auto-reactive cytotoxic T cells that target melanocytes in the basal epidermis. The pathogenic role of CD8^+^ T cells was first reported when the number of patients’ melanocyte antigen-specific CD8^+^ T cells correlated with vitiligo disease activity (Ogg et al., 1998, Palermo et al., 2001), and later confirmed when melanoma patients developed vitiligo after receiving *ex vivo* enriched autologous melanocyte antigen-specific CD8^+^ T cell therapy (Dudley et al., 2002, Yee et al., 2000). To model this process in mice, we adoptively transfer Thy1.1^+^ transgenic PMEL CD8^+^ T cells that recognize melanocyte premelanosome protein (pmel) into Krt14-Kitl* mice that retain epidermal melanocytes to study the effector phase of depigmentation in vitiligo (Harris et al., 2012, Riding et al., 2019). Transferred pmel-specific CD8^+^ T cells (PMELs) identify their target through surveilling peptides presented by the melanocyte major histocompatibility complex class I molecules (MHC-I). After identifying their target in the epidermis, PMELs secrete interferon-gamma (IFNγ), which stimulates the production of C-X-C motif chemokine ligand 9 and 10 (CXCL9 & 10) from surrounding keratinocytes that in turn recruits additional PMELs, expanding the vitiligo lesion (Rashighi et al., 2014, Richmond et al., 2017a). Our mouse model recapitulates several key type I cytokine signatures of human vitiligo lesional skin, including increased CD8^+^ T cell infiltration, up-regulated human leukocyte antigen (HLA) class I molecules on melanocytes (Gellatly et al., 2021, Rashighi et al., 2014), and local CXCL9 concentration as a biomarker of disease activity (Strassner et al., 2017). Hence, this mouse model serves as a valuable tool in the search of better treatments for vitiligo (Azzolino et al., 2021, Richmond et al., 2017b).

The depigmented patches in vitiligo usually develop in a bilateral, symmetrical pattern (Ezzedine et al., 2012). Current treatments include topical calcineurin inhibitors, corticosteroids, and/or narrowband ultraviolet B phototherapy (Rodrigues et al., 2017), while targeted immunotherapy is of significant interest (Rosmarin et al., 2020). We have shown that mice harboring a conditional knockout of signal transducer and activator of transcription (STAT)-1 in keratinocytes blocked the recruitment of PMEL into the skin (Richmond et al., 2017a), however mice harboring type I interferon receptor knockout did not (Riding et al., 2021), indicating that keratinocytes promote T cell recruitment through IFNγ. Thus, preventing IFNγ signaling in keratinocytes may provide a targeted treatment option, a hypothesis consistent with the observed efficacy of topical ruxolitinib in vitiligo (Rosmarin et al., 2020).

Keratinocytes use desmosomes, adherens junctions, and tight junctions to build a tight, integrated epidermal barrier (Sumigray and Lechler, 2015). Desmosomal cadherins, including desmogleins (Dsg) and desmocollins (Dsc), bridge the intermediate filament (keratin) network across cell boundaries, providing mechanical strength to the epidermis. The inverse-gradient expression of Dsg/Dsc isoforms within stratified epithelia is coupled to the differentiation process of keratinocytes – basal layer keratinocytes use Dsg2 along with Dsg3/Dsc2/Dsc3 while the suprabasal layer keratinocytes use more Dsg1/Dsg4/Dsc1 (Garrod et al., 1996). Prior work by Payne et al. described an anti-human Dsg3/Dsg1 single-chain variable fragment (scFv), (D31)12b/6, which bore cross-reactivity to mouse skin but did not cause skin blistering (Payne et al., 2005). Kouno et al. repurposed this scFv for drug delivery to keratinocytes by the name of Px44 and showed a fusion protein that combined the Px44 and tumor necrosis factor–related apoptosis-inducing ligand (Px44-TRAIL) exerted *in vitro* biological activity (Kouno et al., 2013). However, whether the total available Dsg in the skin could support a tissue drug concentration to reach a therapeutic level has not been tested in a disease model *in vivo*. Vitiligo could potentially benefit from Dsg-targeted delivery of IFNγ antagonists as melanocytes reside in the basal layer of epidermis and basal keratinocytes are the largest epidermal source of CXCR3-ligand chemokines in vitiligo patients (Gellatly et al., 2021). Here we fused the Px44 scFv to an IFNγ neutralizing antibody (anti-IFNγ) and determined that this bispecific antibody (BsAb) diminished keratinocyte chemokine production after IFNγ challenge. Contrary to the non-tethered anti-IFNγ antibody, which accumulated in the circulation *in vivo*, the BsAb concentrated locally at the injection site, improving its immunosuppression profile. Finally, our BsAb conferred localized protection to the treated skin, suggesting that tethering an anti-IFNγ antibody to Dsg-expressing keratinocytes can be a viable treatment strategy for vitiligo.

## RESULTS

### Systemic IFNγ neutralization limits vitiligo severity in the mouse model

We recently generated an antigen-pulsed bone marrow-derived dendritic cell (BMDC)-based vitiligo mouse model and reported that disease is dependent on host IFNγ signaling (Riding et al., 2021). To induce vitiligo, irradiated Krt14-Kitl* mice receive co-transfer of Thy1.1^+^ transgenic PMEL CD8^+^ T cells and BMDCs loaded with the pmel cognate peptide. Mice develop more rapid epidermal depigmentation with greater penetrance than disease induced by the recombinant vaccinia virus (rVV). To further validate the pathogenic role of IFNγ in the BMDC model, we treated mice with anti-IFNγ intraperitoneally for 7 weeks after vitiligo induction. IFNγ neutralization led to a reduced disease burden (Figure 1a and 1b), lower skin PMEL number (Figure 1c and S1a), and reduced CXCL9 production by keratinocytes of the vitiligo tail skin (Figure 1d). We also observed significant reduction of CXCL9 and CXCL10 fluorescent reporter proteins in the basal keratinocytes with our rVV model using IFNγR1-deficient REX reporter mice (Figure S2). Consistent with our previous findings (Rashighi et al., 2014, Richmond et al., 2017a, Richmond et al., 2017b), these data demonstrate that the reduction of keratinocyte-derived CXCR3-ligand chemokines correlates with the protective effect of IFNγ blockade, regardless of whether BMDC or rVV is the priming agent for PMEL T cells to induce vitiligo.

**Figure 1.**
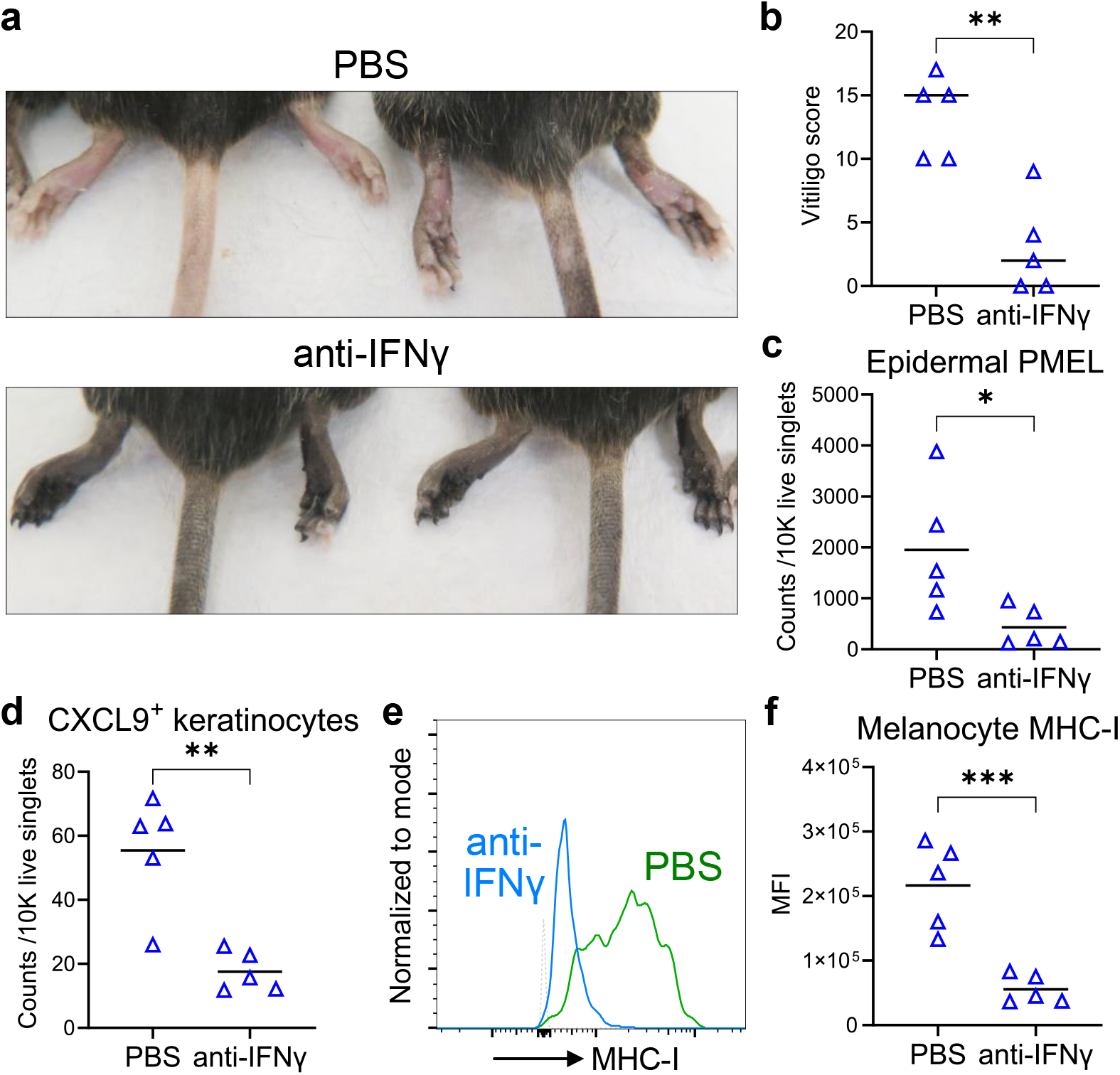
Systemic administration of anti-IFNγ diminishes epidermal inflammation in the BMDC-PMEL vitiligo model. Krt14-Kitl* mice received PBS or 250 ug anti-IFNγ intraperitoneal injections once per week for 7 weeks after BMDC and Thy1.1^+^ PMEL T cell co-transfer. (**a**) Representative image of skin depigmentation in footpads and tail skin compared to anti-IFNγ therapy. (**b**) Vitiligo scores, (**c**) tail epidermal PMEL infiltration, (**d**) CXCL9^+^ tail keratinocyte (gated by CD45^-^, keratin^+^) number, and (**e** and **f**) H-2K/H-2D expression of tail skin melanocytes (gated by CD45^-^, CD117^+^, CD49f^-^, keratin^-^). (**e**) Representative histogram of H-2K/H-2D expression; dotted line indicates the fluorescence minus one (FMO) control for H-2K/H-2D. (**f**) Geometric mean fluorescence intensity (MFI) calculated from **e** for all mice. Data means are indicated, except in **b** which are medians. Data from n=5 mice per group. Student’s *t*-test; \**P* < 0.05, \*\**P* < 0.01, \*\*\**P* < 0.001. All results were reproduced in another independent experiment.

Normal human HLA-A2^+^ melanocytes are lysed by the melanocyte-specific, HLA-A2-restricted CD8^+^ T cell clones isolated from vitiligo patients (Mantovani et al., 2003) and a single-nucleotide polymorphism haplotype in human that increases HLA-A expression is associated with higher vitiligo risk (Hayashi et al., 2016), suggesting that melanocyte MHC-I antigen presentation is important for autoreactive CD8^+^ T cell recognition in vitiligo. Here we found that melanocytes from anti-IFNγ-treated mice had a reduced MHC-I expression (Figure 1e and 1f). Thus, one potential mechanism of protection by the antibody may be through decreasing PMEL recognition of melanocytes. To determine whether anti-IFNγ treatment reduced the magnitude of IFNγ production or altered the phenotype of CD8^+^ PMELs (Sad et al., 1995), we re-stimulated PMELs taken from the skin-draining lymph nodes of treated mice and measured cytokine production by flow cytometry. We did not observe any change in cytokine profile (Figure S1d–S1f), suggesting that systemic anti-IFNγ treatment worked primarily within the skin to reduce keratinocyte production of chemokines as well as autoreactive T cell recruitment to inhibit the development of vitiligo.

### Dsg-IFNγ bispecific antibody preserves the ability to bind both IFNγ and target keratinocytes

Because the suppression of IFNγ-induced immune responses in epidermis correlated with protection of host mice from developing vitiligo, we hypothesized that vitiligo could be locally treated with skin-targeted anti-IFNγ to avoid potential side effects from systemic treatment. To create a skin-targeted BsAb, we cloned the rearranged variable region genes from anti-IFNγ XMG1.2 hybridoma (Cherwinski et al., 1987), and genetically fused the Px44 scFv to the C-terminus of rat IgG1 heavy chain (Figure 2a). The purified BsAb had favorable properties and was a homodimer by SDS-PAGE (Figure 2b and S3a). The purified antibodies only had minute impurity and were non-pyrogenic (Table S1). The BsAb and chimeric anti-Dsg bound to recombinant mouse Dsg3 in ELISA (Figure 2c). In a competition for IFNγ assay, our anti-IFNγ, BsAb, and the commercial XMG1.2 all exhibited comparable affinity to IFNγ (Figure 2d). The BsAb and anti-Dsg both bound to single-cell suspensions from human HaCaT keratinocytes, mouse Pam212 keratinocytes, and mouse tail skin at similar EC_50_ (Figure 2e, S3b, and S3c). To determine whether the BsAb could target the epidermis *in vivo*, biotinylated BsAb or anti-IFNγ were injected into mouse footpads and quantified 24 hrs later by cell surface fluorescently tagged streptavidin on flow cytometry. The BsAb persisted on the basal and suprabasal keratinocytes whereas anti-IFNγ only showed minimal binding (Figure 2f and 2g). This data demonstrates that the BsAb is stable, inherits the dual specificities with similar parental affinity, and can adhere to the epidermis *in vivo*.

**Figure 2.**
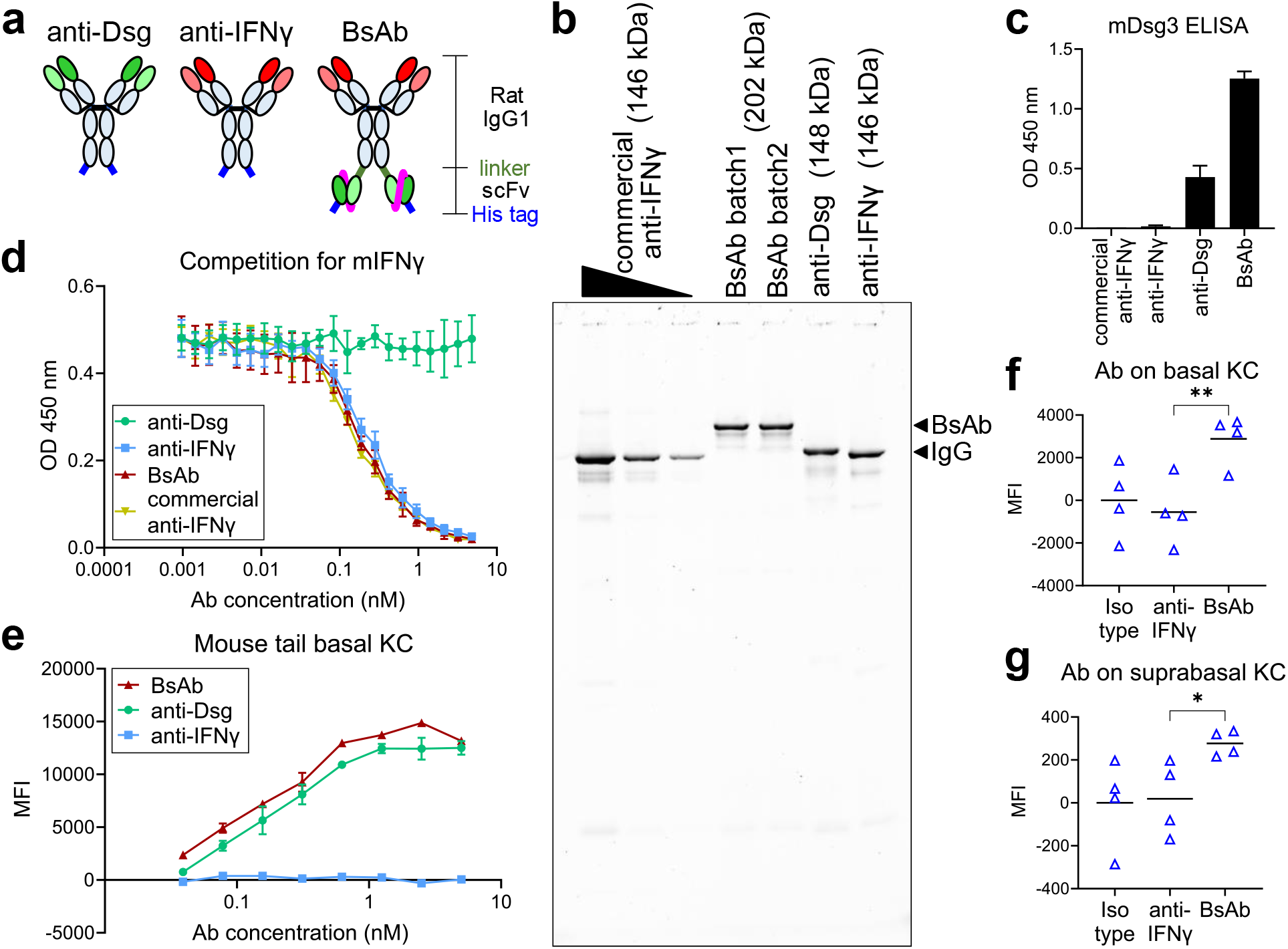
The generation of Dsg-IFNγ bispecific antibody. (**a**) The structure of 6xHis tagged rat IgG1 chimeric anti-Dsg (clone Px44), anti-IFNγ (clone XMG1.2), and the genetically fused BsAb. (**b**) Purified antibodies were subjected to non-reducing SDS-PAGE and Krypton fluorescent protein stain. XMG1.2 antibody from a commercial source served as a benchmark. The theoretical molecular weights of the antibodies are indicated. (**c**) Indirect ELISA against recombinant mouse Dsg3. (**d**) Affinity to IFNγ were evaluated in a competition for recombinant IFNγ ELISA performed on plates coated with the commercial XMG1.2 antibody. Ab, antibody. (**e**) Antibody binding to the mouse tail basal keratinocytes (KC, gated by CD49f^+^, Dsc1^-^, Dsc2/3^+^). (**f** and **g**) Injection site (**f**) CD49f basal and (**g**) CD49f^-^ suprabasal keratinocytes (KC, gated by CD45^-^, keratin^+^) were stained with streptavidin-AF647 to detect surface-bound biotinylated antibodies that had been injected into mouse footpads 24 hrs prior. Data are expressed as mean±SD. One-way ANOVA, Bonferroni post *hoc* test; \**P* < 0.05, \*\**P* < 0.01. All results were reproduced in another independent experiment.

### The BsAb inhibits IFNγ signaling on keratinocytes

Although vitiligo lesional keratinocytes produce the bulk of IFNγ-inducible chemokines, skin-resident immune cells and dermal stromal cells also contribute to the total chemokine production (Richmond et al., 2017a, Richmond et al., 2019, Xu et al., 2021). Considering this, we decided to employ multiple skin models with various cell compositions to assess whether the BsAb could block IFNγ stimulation after binding to keratinocytes. Our first skin model consisted of a confluent layer of HaCaT cells co-cultured with mouse 3T3-J2 fibroblasts. The Dsg-expressing human HaCaT cells responded to human IFNγ stimulation but not to mouse IFNγ (Figure S4a), so we included 3T3-J2 in the culture as detector cells for mouse IFNγ through the production of IFNγ-inducible chemokines (Figure S4b). We treated these cultures with the BsAb, non-tethered anti-IFNγ, or an isotype control antibody for 2 hrs, washed the cultures, and then challenged with mouse IFNγ plus TNFα for 18 hrs. The BsAb pre-treatment substantially inhibited CXCL10 production by 3T3-J2 cells compared to anti-IFNγ or isotype control antibody (Figure 3a). We observed a similar chemokine reduction by the BsAb, albeit to a lesser extent, in another model using only confluent Pam212 cells (Figure S4c and 3b). Lastly, we treated 4 mm mouse tail skin punch biopsy *ex vivo* cultures with the BsAb to evaluate IFNγ neutralization within intact skin *ex vivo*. CXCL9 and CXCL10 production by skin biopsies were IFNγ dose-dependent (Figure S4d). The BsAb pre-treatment of skin biopsies reduced CXCL9 and CXCL10 production compared to biopsies pre-treated with non-tethered anti-IFNγ (Figure 3c). Reduced chemokine production by the BsAb in all three skin models suggests that the BsAb can be tethered to the keratinocyte surface and neutralize IFNγ within the skin.

**Figure 3.**
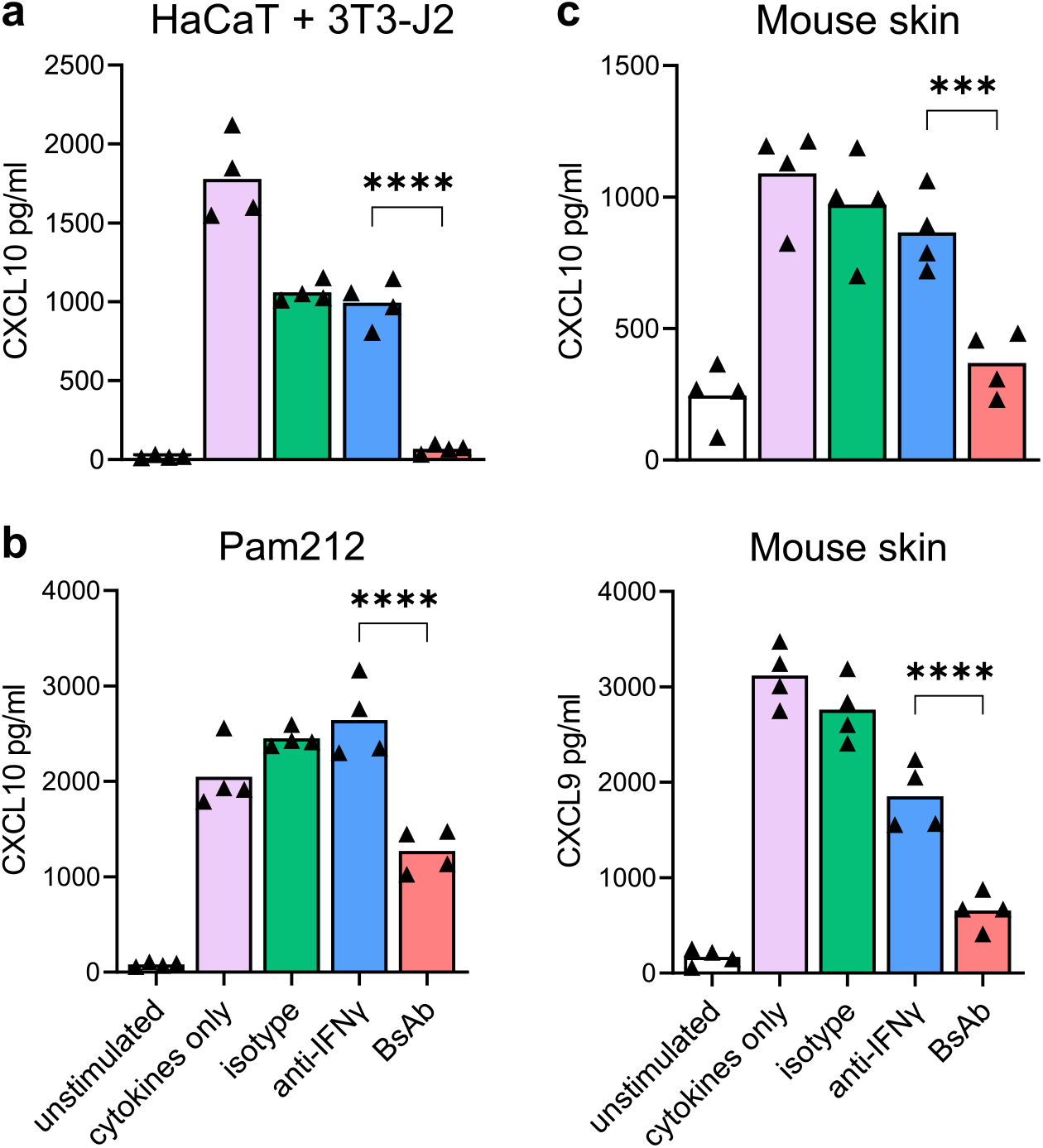
The BsAb protects *in vitro* and *ex vivo* skin cultures from IFNγ challenge. 100% confluent cultures of (**a**) human HaCaT keratinocyte & mouse 3T3-J2 fibroblast lines co-culture, or (**b**) mouse Pam212 keratinocyte line, or (**c**) 4 mm tail skin punch biopsies from treatment-naïve mice were incubated with the antibodies for 2 hrs and washed 3 times to remove unbound antibodies. The skin models were then challenged with IFNγ plus TNFα for 18 hrs and the supernatant was tested for IFNγ-inducible chemokines by ELISA. Data means are indicated. One-way ANOVA, Bonferroni post *hoc* test; \*\*\**P* < 0.001, \*\*\*\**P* < 0.0001. All results were reproduced in another two independent experiments.

### Enhanced skin retention and more rapid elimination of the BsAb lead to reduced systemic IFNγ inhibition

Because the BsAb potentiates immobilized IFNγ-neutralizing activity on keratinocytes, we expected this unique property could synergize with local BsAb administration to reduce the systemic IFNγ inhibition accompanied with treatment. We chose mouse hind footpad skin as the site of testing because the symmetrically depigmented lesions (Figure S5a) allowed us to compare the BsAb-treated and vehicle-treated skin within the same mouse and the Px44 binding capacity of footpad keratinocytes did not change during the development of vitiligo lesions (Figure S5c–S5e). We used iodine-125 labeling to determine the spatial distribution of the BsAb over time. Our ^125^I-labeled anti-IFNγ and BsAb had a similar level of iodination (Table S2). We injected the right footpad of mice with a single dose of 7.1 pmol ^125^I-labeled anti-IFNγ or BsAb, and PBS injected into the left footpad as a control and used single-photon emission computed tomography plus X-ray computed tomography (SPECT/CT) scans to visualize antibody localization.

At 0.5 hrs, both BsAb and non-tethered anti-IFNγ began to diffuse within the injected footpad. The right-side popliteal, iliac, and inguinal lymph nodes exhibited some radioactivity (Figure 4a, Video S1 and S2), indicating that unbound antibodies drained along with extracellular fluid to those lymph nodes. For the first 4 hrs post treatment, both antibodies accumulated in the injection site to a similar level. Long-term retention of the BsAb at the site of injection was clear at 28 hrs and maintained through 48 hrs (Figure 4b). In contrast, non-tethered anti-IFNγ disseminated into the circulation and distal tissues (Figure 4c). The disseminated antibodies constituted most of the whole-body radiation in the anti-IFNγ-treated group, but not in the BsAb-treated group (Figure 4d). The disseminated BsAb signal was primarily detected in urine inside the bladder (Video S4) and induced greater radioactivity signal in the urine compared to anti-IFNγ treatment, suggesting that the BsAb underwent accelerated catabolism, correlating to decreased whole-body radiation following the BsAb treatment (Figure 4d and Table S3). Some BsAb transiently traveled through the circulation and reached distal skin sites to a modest degree, which was less than the level from diffused anti-IFNγ (Figure S6). Consistent with previous studies reporting expression of mouse Dsg3 in various tissues (Culton et al., 2015), we observed weak but focal BsAb signals in the tongue and esophagus (Video S6 and S8), but their intensities were lower than the more dispersed signals from the same head/neck region of the non-tethered anti-IFNγ-treated mice (Video S5 and S7).

**Figure 4.**
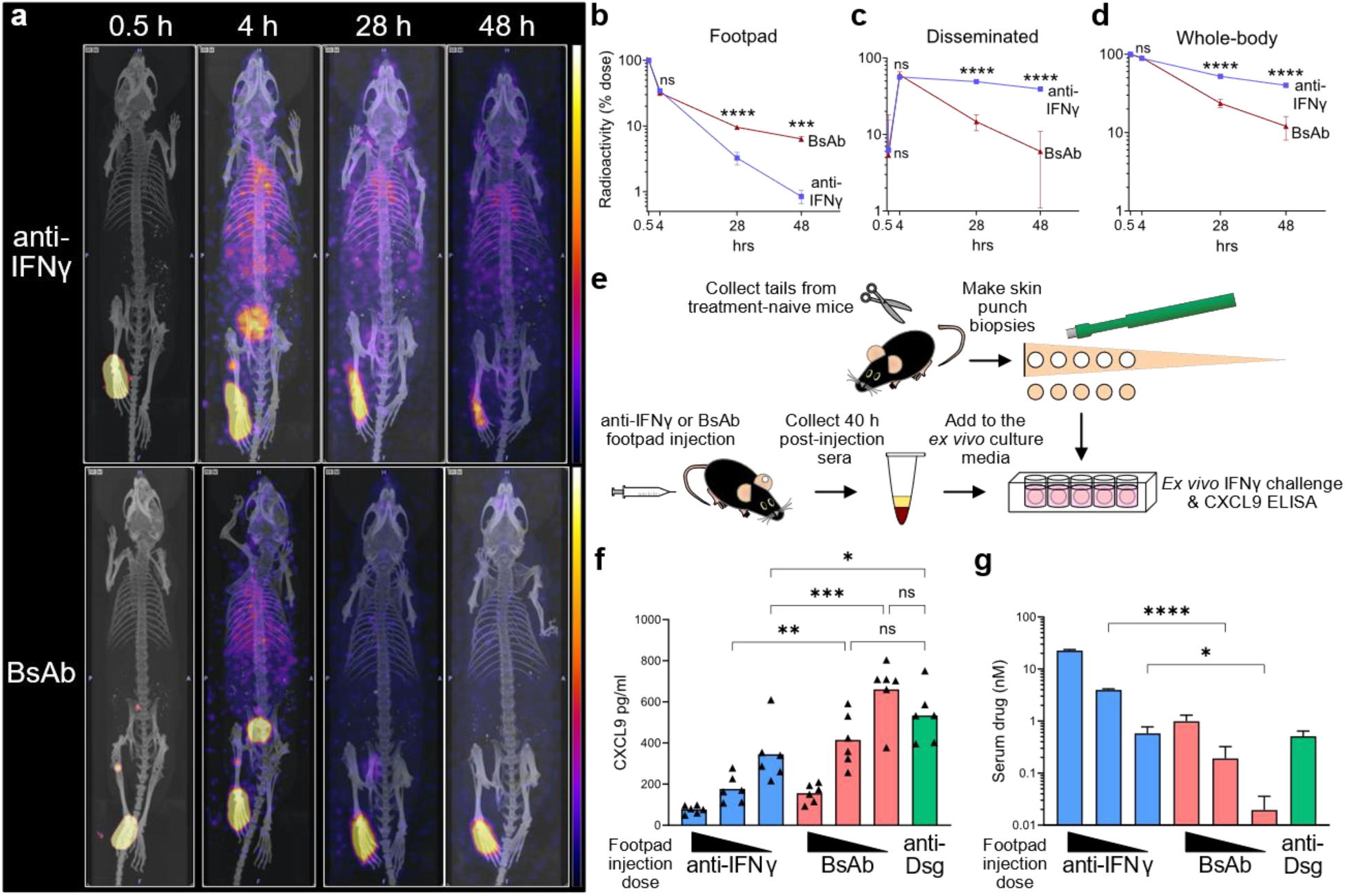
The BsAb has increased local adhesion and reduced systemic IFNγ inhibition than its parental anti-IFNγ. Krt14-Kitl* mice received 7.1 pmol of ^125^I-labeled BsAb or anti-IFNγ injected in the right footpad and PBS in the left. (**a**) Representative SPECT/CT image of one mouse from each group are shown. (**b**) Radioactivity quantification of SPECT signal from the right footpad; data was normalized to the measurement taken at 0.5 hrs. (**c**) Disseminated radioactivity was calculated by subtracting radioactivity of right footpad from (**d**) the whole-body radioactivity as measured by a radioisotope calibrator. Data are expressed as mean±SD from n=3 mice per group, compiled from two independent experiments. (**e**) Schematic of measuring the disseminated IFNγ-neutralizing activity in serum. Footpads were injected with the BsAb or anti-IFNγ at 28.3, 7.1, or 1.8 pmol; anti-Dsg was given a single 28.3 pmol dose. (**f**) CXCL9 production by the IFNγ-stimulated skin in the presence of 5% mouse sera from the antibody recipients. (**g**) ELISA measurement of the concentration of injected antibodies in serum. Data are expressed as mean±SD. Two-way ANOVA, Bonferroni post *hoc* test (**b**-**d**) or one-way ANOVA, Holm-Sidak post *hoc* test (**f** and **g**); ns, not significant, \**P* < 0.05, \*\**P* < 0.01, \*\*\**P* < 0.001, \*\*\*\**P* < 0.0001. Results in (**f** and **g**) were reproduced in another independent experiment.

We observed a significant difference between the distribution patterns of anti-IFNγ and our BsAb in that anti-IFNγ accumulated in the circulation, whereas the BsAb was retained at the injection site. We next tested whether the antibody diffused from serum would functionally impair the IFNγ responsiveness of vitiligo-unaffected distal skin sites as an off-tissue effect by the drug. To do this we used tail skin punch biopsies from naïve mice to model uninvolved skin that does not require treatment. We stimulated skin biopsies with IFNγ in the presence of 5% serum from mice that had received a single dose of anti-Dsg, anti-IFNγ, or BsAb by footpad injection (Figure 4e). As low as 1.8 pmol anti-IFNγ footpad injection resulted in sera capable of reducing naive skin CXCL9 production compared to the sera from anti-Dsg control; whereas sera from mice treated with 7.1 pmol BsAb did not significantly decrease the amount of CXCL9 in skin biopsies (Figure 4f). This suggests that the BsAb causes less systemic immunosuppression and off-tissue effect than non-tethered anti-IFNγ. ELISA quantification of the sera revealed 20-fold more anti-IFNγ in molar concentration than the BsAb in each injection dose tested (Figure 4g). These results demonstrate that the BsAb increases local drug availability and limits systemic exposure to the body. Since we observed that the BsAb is eliminated from the circulation more rapidly, its systemic IFNγ inhibition is significantly lower than that by local administration of a non-targeted anti-IFNγ.

### Dsg-targeting strategy confers selective protection in treated skin

After characterizing the skin-retention of the BsAb *in vivo*, we determined whether the BsAb local administration would be sufficient to protect skin depigmentation locally in our BMDC mouse vitiligo model. We injected the right footpad with antibodies and treated with the same volume of PBS to left footpad every other day throughout the experiment following vitiligo induction. We photographed the footpad each week and quantified the percentage of pigmented area using ImageJ. Anti-Dsg-treated mice showed nearly symmetrical depigmentation in both footpads (Figure 5a). The average difference in pigmented area between the two footpads (right minus left footpad, referred to as “Δ”) in the anti-IFNγ group was increased but not statistically significant because depigmentation was concurrently slowed down in both feet (Figure S7a and S7b). The small elevation in Δ pigmented area likely came from the residual anti-IFNγ at the injection site observed 48 hours post injection (Figure 4a). Only the BsAb-treated footpads showed a significant Δ pigmented area (Figure 5a and 5b), indicating that the protective effect of the BsAb was confined to the injection site (Figure S7a) and extended very little to the contralateral footpad or other body locations (Figure S7b and S7c), which correlates with the low systemic IFNγ inhibition property of the BsAb (Figure 4).

**Figure 5.**
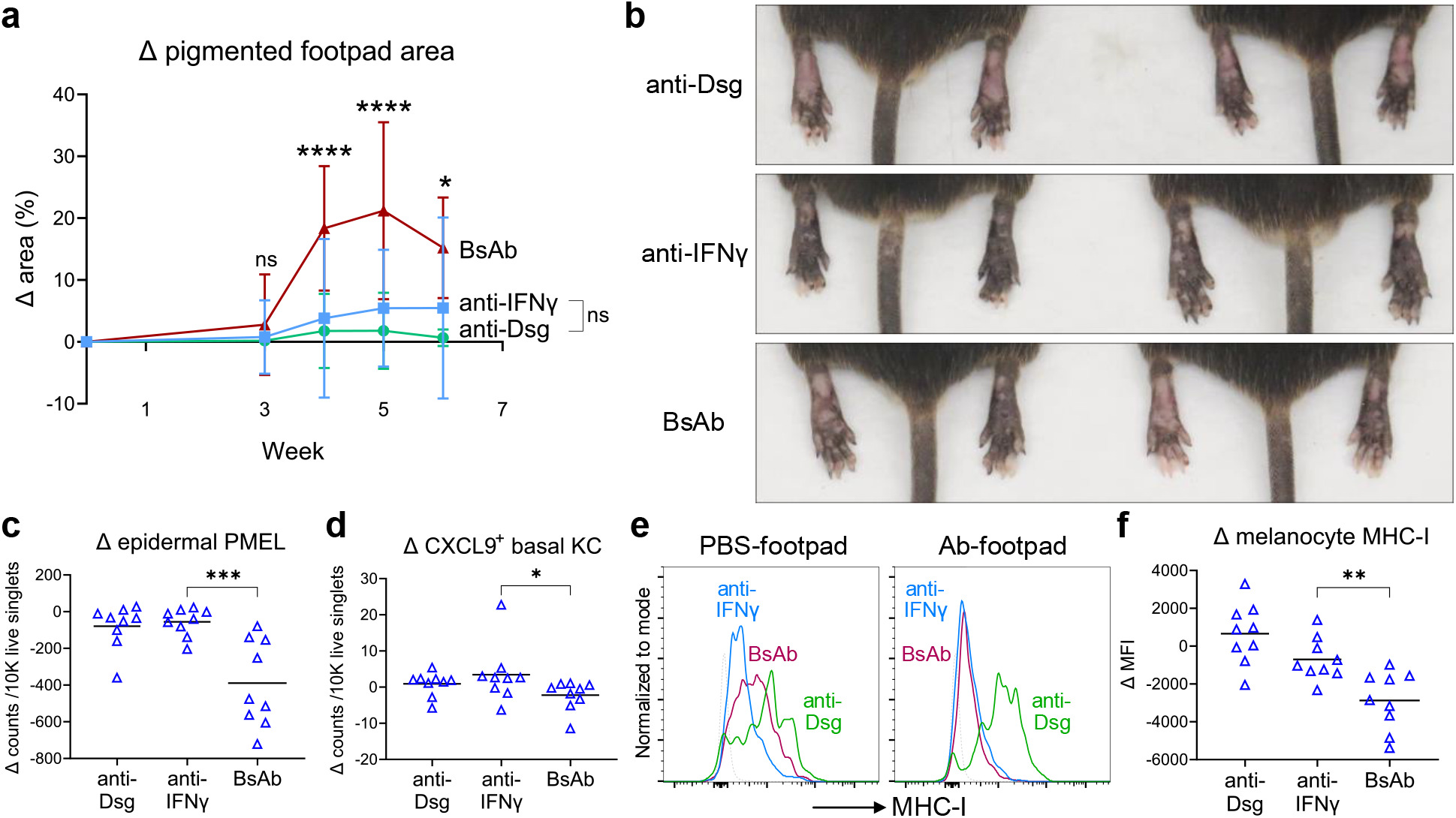
Dsg targeting-dependent localized skin pigment protection by the BsAb. Krt14-Kitl* mice received 7.1 pmol antibody to the right hind footpad and PBS to the left every two days following vitiligo induction. (**a**) The percentage of pigmented area in each footpad was quantified by ImageJ analysis of mouse photographs and the mathematical difference (right minus left footpad, referred to as “Δ”) of each mouse was plotted. Data were compiled from two independent experiments. BsAb, n=14 mice; anti-IFNγ, n=14 mice; anti-Dsg, n=9 mice. Treatment differences compared to the group receiving anti-IFNγ at each time point were tested by two-way ANOVA with Bonferroni post *hoc* test. (**b**) Representative photos of each treatment group after 5 weeks, (**c**) Δ footpad epidermal Thy1.1^+^ PMEL T cell infiltration, (**d**) Δ CXCL9^+^ basal keratinocytes (KC, gated by CD49f^+^, keratin^+^) number, (**e**) representative H-2K/H-2D expression of melanocytes from footpad skin (gated by CD45^-^, CD117^+^, CD49f^-^, keratin^-^), FMO indicated by the dotted line, and (**f**) Δ melanocyte H-2K/H-2D MFI at week 5 of treatment. Data are expressed as mean±SD. One-way ANOVA with Bonferroni post *hoc* test (**c**, **d**, **f**); ns, not significant, \**P* < 0.05, \*\**P* < 0.01, \*\*\**P* < 0.001, \*\*\*\**P* < 0.0001. All results were reproduced in two independent experiments.

Consistent with this asymmetrical depigmentation, the BsAb exclusively decreased the PMEL infiltration into right footpad dermis and epidermis (Figure S8a and 5c). The differential PMEL recruitment correlated closely with the basal keratinocyte trafficking-related chemokine CXCL9 production pattern (Figure 5d) but less strongly with the dermal stromal cell CXCL9 production pattern (Figure S8b). Since our SPECT/CT scan revealed the drainage of unbound antibodies into the right-side lymph nodes during footpad injection (Figure 4a), we asked whether the PMELs in those popliteal lymph nodes had impaired migration. We did not find any difference in PMELs from the draining versus opposite lymph nodes in terms of PMEL engraftment, cognate peptide stimulated IFNγ production, or CXCR3 receptor expression (Figure S9b–S9e). Thus, the BsAb-induced differential PMEL infiltration was more likely due to reduced chemokine production in the skin, rather than any desensitizing predisposition in the draining lymph nodes.

Lastly, analysis of melanocytes at the injection site indicated that the BsAb and anti-IFNγ treatment regimen had similar potency to decrease melanocyte MHC-I expression level (Figure 5e) and possibly evade PMEL recognition. However, unlike the systemic effect of anti-IFNγ, the BsAb only had a minor influence on melanocytes of the opposite footpad (Figure 5f). Together, these results demonstrate that a keratinocyte-tethering strategy can functionally preserve the ability of anti-IFNγ to limit depigmentation and deliver a therapeutic effect in a location-precise manner that cannot be achieved by local injection of the non-tethered antibody alone.

## DISCUSSION

The function of a novel BsAb in a vitiligo mouse model suggests that active skin retention can significantly enhance the localized therapeutic effect of a locally delivered biologic. The skin-tethering antibody clone Px44 was one of the non-blistering clones isolated from a patient with pemphigus vulgaris, and it recognizes a calcium-insensitive epitope in the extracellular cadherin domain 1 (EC1) in Dsg3 rather than the calcium-stabilized epitopes targeted by most pathogenic clones (Cho et al., 2014, Payne et al., 2005). Consistent with these reports, footpads treated with anti-Dsg or the BsAb did not exhibit any epidermal erythema, which is the primary skin manifestation in mouse pemphigus models (Lotti et al., 2019). We did not introduce the Fc-function-null mutations into our antibodies because of the location of Px44 epitope within Dsg3. The isolation of anti-Dsg3 antibody clones that can provoke strong antibody-dependent cellular cytotoxicity (ADCC) concluded that those epitopes were not in EC1 (Funahashi et al., 2018), probably due to EC1 being distant from the cell membrane and not amenable to immune synapse formation or transfer of cytotoxic granules (Cleary et al., 2017, Funahashi et al., 2018). In addition, we anticipated the risk of stressing melanocytes by Dsg-targeted drug delivery would be low since melanocytes utilize α6β1 integrins and E-cadherin to adhere themselves to the basement membrane and keratinocytes, respectively (Hara et al., 1994, Tang et al., 1994). We did not find the binding of Px44 on melanocytes and as expected, the average pigmented area in anti-Dsg-treated footpads was not lower than their contralateral side, or the average of untreated vitiligo mice.

Tracking ^125^I-labeled BsAb confirmed keratinocyte-tethering and explained the differential footpad depigmentation in the vitiligo mouse model. Nonetheless, the slightly higher (statistically insignificant) average pigmented area in the anti-IFNγ-treated right footpads, compared to the BsAb-treated footpads, may seem surprising given that the BsAb retained in footpads longer than anti-IFNγ did. We believe it was because those mice in our SPECT/CT study had only received a single dose of antibody and the biodistribution data did not capture the accumulation of anti-IFNγ in serum after repetitive dosing like in our vitiligo model. In fact, the serum anti-IFNγ to BsAb molar concentration ratio increased from 20-fold after the 1^st^ dose to more than 400-fold after 3 weeks of antibody treatment (data not shown). Hence, unless the two compared tissue-targeted/non-targeted compounds have similar serum half-lives like in these studies (Pemmari et al., 2020, Sum et al., 2021), any enhanced therapeutic effects of our BsAb due to concentrating locally in the treated skin area would have been masked by the increasing anti-IFNγ reservoir of blood diffusing back to the tissue.

The imbalanced half-lives of anti-IFNγ and the BsAb originate from the rapid BsAb elimination *in vivo*. At least five factors affect the elimination rate of immunoglobulins (Keizer et al., 2010). Briefly, FcγR binding-mediated endolysosomal degradation by phagocytes in the reticuloendothelial system, neonatal Fc receptor (FcRn) binding-mediated recycling from endocytosed vesicles, molecular size-associated filterability at the glomerulus, target antigen-binding mediated endolysosomal degradation, and patient acquired anti-drug immune responses. Although both BsAb and anti-IFNγ are large (> albumin, 69 kDa) and bear the same Fc sequence, we did not test how the first three factors might contribute to differences in half life. Nevertheless, binding to the target Dsg^+^ keratinocytes and epithelial cells would promote depletion of the BsAb from the circulation. Desmosomes are also dynamic structures that undergo cycles of assembly and turnover to adapt to tissue remodeling (Nekrasova and Green, 2013). A pulse-chase study demonstrated that the biotinylated Dsg3 on the surface membrane of confluently cultured normal human keratinocytes became undetectable 6-24 hrs after initial biotin labeling (Calkins et al., 2006), which may reflect the fate of the BsAb after binding to Dsg3. Furthermore, we have detected anti-rat IgG1 antibodies in the sera of mice receiving anti-IFNγ for five weeks (data not shown). It is not surprising that the BsAb might carry more immunogenicity than anti-IFNγ due to the human-derived Px44 scFvs at the C-terminus of the BsAb. As a result, the BsAb serum half-life may further decrease over time. Yet these pharmacokinetic differences between anti-IFNγ and the BsAb provide the foundation for our observed divergence of systemic versus localized treatment effects.

To further refine and optimize the BsAb kinetics and treatment profile, it may be prudent to further shorten the serum half-life by removing the Fc region from both molecules in the future. This concept has been employed by ranibizumab, FDA-approved for intravitreal injections to treat age-related macular degeneration (AMD). Instead of a whole anti-VEGF IgG molecule (bevacizumab), ranibizumab is a fragment antigen-binding (Fab) variant derived from bevacizumab, designed for better retinal penetration and reduced systemic anti-VEGF exposure (Ferrara et al., 2006). Compared to the off-label use of bevacizumab in AMD, ranibizumab showed equivalent therapeutic effect and decreased systemic adverse events rates in several meta-analyses (Plyukhova et al., 2020, Schmucker et al., 2012), though not all these differences can be attributed to the Fab versus IgG format.

Another important factor to consider is whether localizing anti-cytokine function via bispecific antibodies would create a local depot of cytokine within the tissue, thereby prolonging an inflammatory state. The outcome may depend on the parental clone chosen for anti-cytokine function. For example, certain anti-mouse IL-2 clones like S4B6 and JES6-1 can extend IL-2 half-life *in vivo* by forming immune complexes (Létourneau et al., 2010). Either injecting S4B6 alone or combining with IL-2 preferentially expands IL-2Rβ^hi^ CD8^+^ memory T cells because the S4B6:IL-2 complex structurally mimics the IL-2Rα:IL-2 complex and can form a more stable interaction with IL-2Rβ:IL-2Rγ heterodimer. JES6-1 must combine with exogenous IL-2 before injection to show a T cell expansion effect and the closely matched affinities (K_d_, JES6-1:IL-2 = 8 nM; IL-2Rα:IL-2 = 15 nM) allow predominantly IL-2Rα^hi^ CD4^+^ T cells to displace JES6-1 by IL-2Rα and utilize IL-2 (Boyman et al., 2006, Spangler et al., 2015). Unlike the stimulatory effect by S4B6, administrating XMG1.2 in our vitiligo mouse model suppresses IFNγ-induced MHC-I and chemokine production in the skin. Moreover, XMG1.2 affinity to IFNγ is about 645-fold higher than that of the IFNγ receptor (Basu et al., 1988, Merli et al., 2020). Thus, our BsAb is unlikely to release bound IFNγ before becoming degraded by the Dsg^+^ keratinocytes. Other evidence supports this as well: TNFα and TGFβ neutralizing antibodies equipped with a second targeting moiety against the extracellular matrix in joint, lung, or kidney have shown enhanced treatment efficacy (Katsumata et al., 2019a, Katsumata et al., 2019b, McGaraughty et al., 2017), supporting the hypothesis that prolonging an inflammatory state is not a generalized phenomenon in tissue targeting approaches.

Advances in polymeric biomaterial-mediated drug release also aim to achieve a localized treatment effect. These locally applied systems, including nanotubes, hydrogels, and polymeric scaffolds, gradually release the drug into tissue to reduce dosing frequency, however, can still have systemic drug exposure depending on the cargo pharmacokinetics (Bentley and Little, 2021). Cytotoxic T-lymphocyte associated protein 4-Fc (CTLA4-Fc) immobilized within a matrix and inserted at the transplantation site of a mouse islet allograft maintained the serum CTLA4-Fc at a low and steady level for 8 weeks, compared to the drug level surge that lasted for 2 weeks without using the matrix (Zhang et al., 2016). Our Dsg-tethering strategy may conceptually complement controlled drug release systems and promote both region-specific efficacy and long-acting therapy. In summary, we have used a mouse vitiligo model to validate Dsg as a target for converting current and future biologics into skin-localized therapeutics. This strategy synergizes with the direct treatment of skin lesions to limit off-tissue effect *in vivo*.

## MATERIALS AND METHODS

### Mice

All mice were bred and housed in pathogen-free facilities at UMMS. Procedures were approved by the UMMS Institutional Animal Care and Use Committee in accordance with the NIH Guide for the Care and Use of Laboratory Animals. The following mouse strains used in this study are available from The Jackson Laboratory: KRT14-Kitl*4XTG2Bjl (Krt14-Kitl*) mice (stock no. 009687); Thy1.1^+^ PMEL TCR transgenic mice (stock no. 005023); Thy1.1^+^ PMEL TCR transgenic mice were bred to DPE^GFP^ mice (Mrass et al., 2006) (provided by Dr. Ulrich von Andrian, Harvard Medical School) to generate GFP^+^ PMEL TCR transgenic mice; IFNγR1-KO mice (stock no. 003288), REX3 mice (Groom et al., 2012) (provided by Dr. Andrew Luster, Massachusetts General Hospital), and Krt14-Kitl* mice were bred together to generate REX3^+^, IFNγR1-KO, Krt14-Kitl* recipient mice for vitiligo induction. All mice were on C57BL/6J background or had been backcrossed to C57BL/6J background for more than ten generations.

### Vitiligo induction

BMDC-induced vitiligo was adapted from the previously described method (Riding et al., 2021). Bone marrow cells taken from the femurs and tibia of 8–16-week-old C57BL/6J mice after RBC lysis (MilliporeSigma) were cultured in BMDC differentiation medium [RPMI-1640 (MilliporeSigma, R8758), 10% FBS (MilliporeSigma), 100 U/ml penicillin-streptomycin, 20 ng/ml mouse GM-CSF (PeproTech), 10 ng/ml mouse IL-4 (PeproTech), 50 uM 2-mercaptoethanol] at 0.8×10^6^ cells/ml. The medium was changed on day 3 and 6. 8–16-week-old Krt14-Kitl* recipient mice received 500 rad sublethal irradiation on day 7. On day 8, BMDC were pulsed with 10 uM human gp100_25-33_ peptide (GenScript) in Opti-MEM medium (ThermoFisher) in 37°C, 5% CO2 for 3 hrs. The expression of MHC-II, CD11c, CD11b, CD86 were also evaluated by flow cytometry. PMEL T cells were isolated from the splenocytes of the Thy1.1^+^ PMEL TCR transgenic mice by CD8a^+^ negative selection (Miltenyi Biotec). Each irradiated Krt14-Kitl* mouse received 1×10^6^ cells of PMEL plus 1.8×10^6^ cells of the peptide-pulsed MHC-II^+^, CD11c^+^ BMDC through retro-orbital injection on day 8, and 600 IU mouse IL-2 (R & D Systems) by intraperitoneal injection on day 9, 10, and 11. The cell donors and recipients were gender-matched.

Recombinant vaccina virus-induced vitiligo was performed as previously described (Harris et al., 2012). Each 500 rad irradiated Krt14-Kitl* recipient mouse received 1×10^6^ cells of CD8a^+^ negative selection-enriched GFP^+^ PMEL through retro-orbital injection and 1×10^6^ pfu rVV-hPMEL virus (provided by Dr. Nicholas Restifo, NIH) through intraperitoneal injection.

### Bispecific antibody production

The sequence of anti-IFNγ was cloned from XMG1.2 hybridoma (ATCC) using SMARTer RACE 5’/3’ kit (Takara Bio). The 6xHis tagged anti-Dsg, anti-IFNγ, and BsAb were expressed by transient transfection in ExpiCHO system (ThermoFisher), followed by HisPur cobalt column (ThermoFisher) purification, and buffer exchange during the concentrating step using Amicon Ultra 100K device (MilliporeSigma). The composition of the purified antibodies was verified by NuPAGE gradient SDS-PAGE (ThermoFisher) in both reducing and non-reducing conditions plus Krypton fluorescent whole protein staining (ThermoFisher). A commercially purchased XMG1.2 (BioXCell) was used as a standard in the SDS-PAGE and NanoDrop protein quantification by individual extinction coefficient (ThermoFisher) assays to determine the concentrations of the purified antibodies. The endotoxin levels of the purified antibodies were measured by Chromogenic Endotoxin Quant Kit (ThermoFisher) according to the manufacturer’s instructions.

### Competition for mouse IFNγ ELISA

To rank the affinity to IFNγ of purified antibodies, 100 pg recombinant mouse IFNγ (R & D Systems) was mixed with a concentration gradient of the purified antibodies in wells that had been coated with a commercial anti-IFNγ (clone XMG1.2, BioXCell). After incubation and washing out the unbound antibody-IFNγ complexes, the IFNγ that had been captured by the plate-bound XMG1.2 was detected by the biotinylated rabbit anti-mouse IFNγ (PeproTech), streptavidin-HRP (Abcam), and TMB substrate (BioLegend). After adding 2N hydrochloric acid solution to stop the enzymatic reaction, the absorbance at 450 nm of each well was read by a SpectraMax M5 microplate reader (Molecular Devices).

### Skin IFNγ challenge assays

Pam212 and HaCaT were grown till 100% confluence at least 24 hrs before the experiment to enable desmosome formation. 3T3-J2 mouse fibroblasts were added to the confluent HaCaT culture 24 hrs before the experiment to reach 20% confluence on the next day. 4 mm tail skin punch biopsies taken from 8-12-week-old treatment naïve C57BL/6J mice were cultured in the keratinocyte growth medium described in (Barreca et al., 1992), except without hydrocortisone. The skin models were incubated with the culture media containing 14 nM of anti-IFNγ, BsAb, or a commercial rat IgG1 (SouthernBiotech) for 2 hrs in 37°C, 5% CO2 and then subjected to three washes with media to remove unbound antibodies. Skin models were then challenged with 0.2 ng/ml mouse IFNγ and 10 ng/ml mouse TNFα (R & D Systems) for 18 hrs. The supernatants from the mouse tail skin punch biopsies were collected for both mouse CXCL9 and CXCL10 ELISA (Abcam) measurements. Because the CXCL9 levels made by 3T3-J2 and Pam212 cells were relatively close to the detection limit of the ELISA kit (Fig S4b and S4c), only CXCL10 ELISA was performed on the supernatants from these two skin models due to assay sensitivity.

### Mouse Dsg binding assays

For ELISA, 10 nM of purified anti-Dsg, anti-IFNγ, BsAb, or commercial XMG1.2 (BioXCell) was added to the ELISA plates that had been coated with 2 ug/ml recombinant mouse Dsg3 protein extracellular domains (MyBioSource, baculovirus expressed). The binding complexes were detected by HRP goat anti-rat IgG (BioLegend) and TMB substrate.

For Dsg^+^ keratinocyte binding curves, Pam212 (provided by Dr. Wendy Havran, The Scripps Research Institute) and HaCaT (AddexBio) keratinocyte lines were grown in 37°C, 5% CO2 in DMEM (CORNING, 10-013-CV) +10% FBS + 100 U/ml penicillin-streptomycin, until 100% confluence at least 24 hrs before the experiment to enable desmosome formation. Mouse tail epidermis was separated from dermis by 5 mg/ml Dispase II (Roche) digestion in 37°C for 1 hr. The monolayer culture/epidermis was mechanically dissociated by rubbing through mesh into single-cell suspensions and then incubated with mouse Fc blocker (2.4G2, BioXCell) and the purified antibodies. The binding complexes were detected by Alexa Fluor 647 goat anti-rat IgG (BioLegend) on MACSQuant (Miltenyi) or Aurora flow cytometry (Cytek).

For profiling the Px44 binding capacity of footpad keratinocytes and melanocytes, epidermal single-cell suspensions were prepared from the hind footpad skin of age-matched naïve Krt14-Kitl* mice and 10-week-post-induction vitiligo mice. The cells were stained with the biotinylated anti-Dsg (clone Px44) or a biotinylated rat IgG1 (clone HRPN), followed by streptavidin-Alexa Fluor 647 (BioLegend) detection and flow cytometry analysis.

### Antibody biodistribution characterization

For characterization of footpad keratinocyte targeting, a rat IgG1 isotype antibody (BioXCell), and purified anti-IFNγ and BsAb were biotinylated by EZ-Link Sulfo-NHS-SS-Biotin (ThermoFisher) and buffer exchanged by Zeba Spin Desalting Columns (7K MWCO, ThermoFisher) according to the manufacturer’s instructions. 10 ul, 2.8 uM of the biotinylated isotype antibody, anti-IFNγ, or BsAb was injected to Krt14-Kitl* mouse right hind footpads and 10 ul PBS to the left hind footpads. The epidermal single-cell suspensions of the right footpads were prepared 24 hrs later and subjected to streptavidin-Alexa Fluor 647 cell surface staining for detection of the biotinylated antibodies.

For characterization of systemic distribution, purified anti-IFNγ and BsAb were radiolabeled with ^125^I-radionuclide (Perkin Elmer) using Iodogen reaction and then purified by P-6 spin column (Bio-Rad) according to the manufacturer’s instructions. 10 ul, 0.7 uM of the iodinated anti-IFNγ, or BsAb was injected to Krt14-Kitl* mouse right hind footpads and 10 ul PBS to the left hind footpads. At the 0.5, 4, 28, and 48 hrs-post injection timepoints, the mouse whole-body radioactivity was measured by CRC-7 radioisotope calibrator (Capintec) and 3D SPECT/CT imaging was performed on a VECTor^6^/CT system (MILabs). After data reconstruction, the volume-of-interest (VOI) analysis on footpads and video representation of the SPECT images were carried out by PMOD 4.201 (PMOD Technologies LLC).

### Testing of vitiligo prevention

To test the effect of systemic vitiligo prevention by anti-IFNγ, mice received weekly intraperitoneal injection of 250 ug anti-IFNγ from a commercial source (BioXCell, clone XMG1.2) or PBS for 7 weeks following BMDC-vitiligo induction. The vitiligo disease scores were graded by a trained investigator who was blinded to the experimental setup. The scale of the scoring point system was described previously (Riding et al., 2019).

To test the effect of localized vitiligo prevention by the BsAb, mice received 10 ul, 0.7 uM of purified anti-Dsg, anti-IFNγ, or BsAb to the right hind footpads and 10 ul PBS to the left hind footpads every other day until the study endpoint. In addition to the vitiligo disease scores, the pigmented footpad areas were also recorded weekly using photographs and quantified by ImageJ version 1.52a (NIH). The toes were excluded from the pigmented area quantification due to high frequency of unpigmented birthmarks.

### Testing of systemic IFNγ inhibition by serum

8–12-week-old C57BL/6J mice received 10 uL of 2.8 uM, 0.7 uM, or 0.2 uM of anti-IFNγ, BsAb, or anti-Dsg in the right footpads and 10 uL PBS for the left footpads. Sera were collected at 40 hrs-post-injection. For ELISA quantification of the serum concentration of the injected antibodies, the sera were applied to 3 ug/ml goat anti-rat IgG (BioLegend), or 1% BSA (ThermoFisher) coated ELISA plates. The pre-quantified anti-IFNγ and BsAb were used as concentration standards. The captured antibodies were detected by HRP goat anti-rat IgG (BioLegend) and subsequent TMB substrate. Readings from the BSA-coated plates were used as the background for individual serum samples. For testing the IFNγ neutralization ability of the sera, 4 mm tail skin punch biopsies taken from the treatment-naïve C57BL/6J mice were cultured in the keratinocyte growth medium. Because a typical IgG drug would establish about 7% of its plasma concentration in mouse skin interstitial space (Chang et al., 2021), the indicated mouse sera were added to the culture media at 5% concentration. The cultures were stimulated with 0.5 ng/ml mouse IFNγ, and 10 ng/ml mouse TNFα (R & D Systems) for 18 hrs. The supernatants were collected for CXCL9 ELISA (Abcam).

### Flow cytometry

At the endpoint of vitiligo prevention mouse experiments, the spleens, dermis, epidermis, and skin-draining lymph nodes were harvested for flow cytometry analyses. Detailed tissue processing procedures are described elsewhere (Riding et al., 2019). Single-cell suspensions of lymph nodes, dermis, and epidermis were cultured overnight in complete RPMI-1640 medium containing 10 uM human gp100_25-33_ peptide. 5 ug/ml Brefeldin A was supplemented 4 hrs before subsequent Fc blocking (2.4G2, BioXCell), surface antibody staining, and Zombie Aqua cell viability staining (BioLegend). The cells were fixed and permeabilized by Cytofix/Cytoperm (BD) before intracellular antigen staining. The stained samples were measured by Aurora flow cytometry (Cytek) and analyzed by FlowJo version 10.8.0 (FlowJo, LLC). Cell population gates were drawn by the isotype antibody staining or FMO control.

Antibodies (clone) purchased from BioLegend: Thy1.1 (OX-7), CD117 (2B8), CD49f (GoH3), H-2K^b^/H-2D^b^ (28-8-6), CXCR3 (CXCR3-173), CD8 (YTS156.7.7), keratin (C-11), IFNγ (XMG1.2), IL-17A (TC11-18H10.1), CXCL9 (MIG-2F5.5), CD11b (M1/70), CD86 (GL-1), I-A/I-E (M5/114.15.2), CD11c (N418). Purchased from Santa Cruz: desmocollin1 (A-4), desmocollin2/3 (7G6). Purchased from NOVUS: IL-4 (11B11). Purchased from BD: CD45 (30-F11).

### Statistical analyses

The statistical evaluations of each experiment were carried out by GraphPad Prism version 9.3.0 (GraphPad Software). The statistical type of each hypothesis testing is indicated in the figure legend. *P* values below 0.05 were considered significant. The significance level symbols used here include: \**P*<0.05, \*\**P*<0.01, \*\*\**P*<0.001, \*\*\*\**P*<0.0001. ns, not significant, *P*>0.05.

## ACKNOWLEDGEMENTS

This work was funded by the Dermatology Foundation and American Skin Association, Hartford Foundation Vitiligo Grant and other philanthropic gifts, and NIH R01 AR069114 (to J.E.H.). The authors thank Dr. Ann Rothstein and Dr. Eric Kercher for their critical comments on this manuscript.

---

**Fig. S1.**
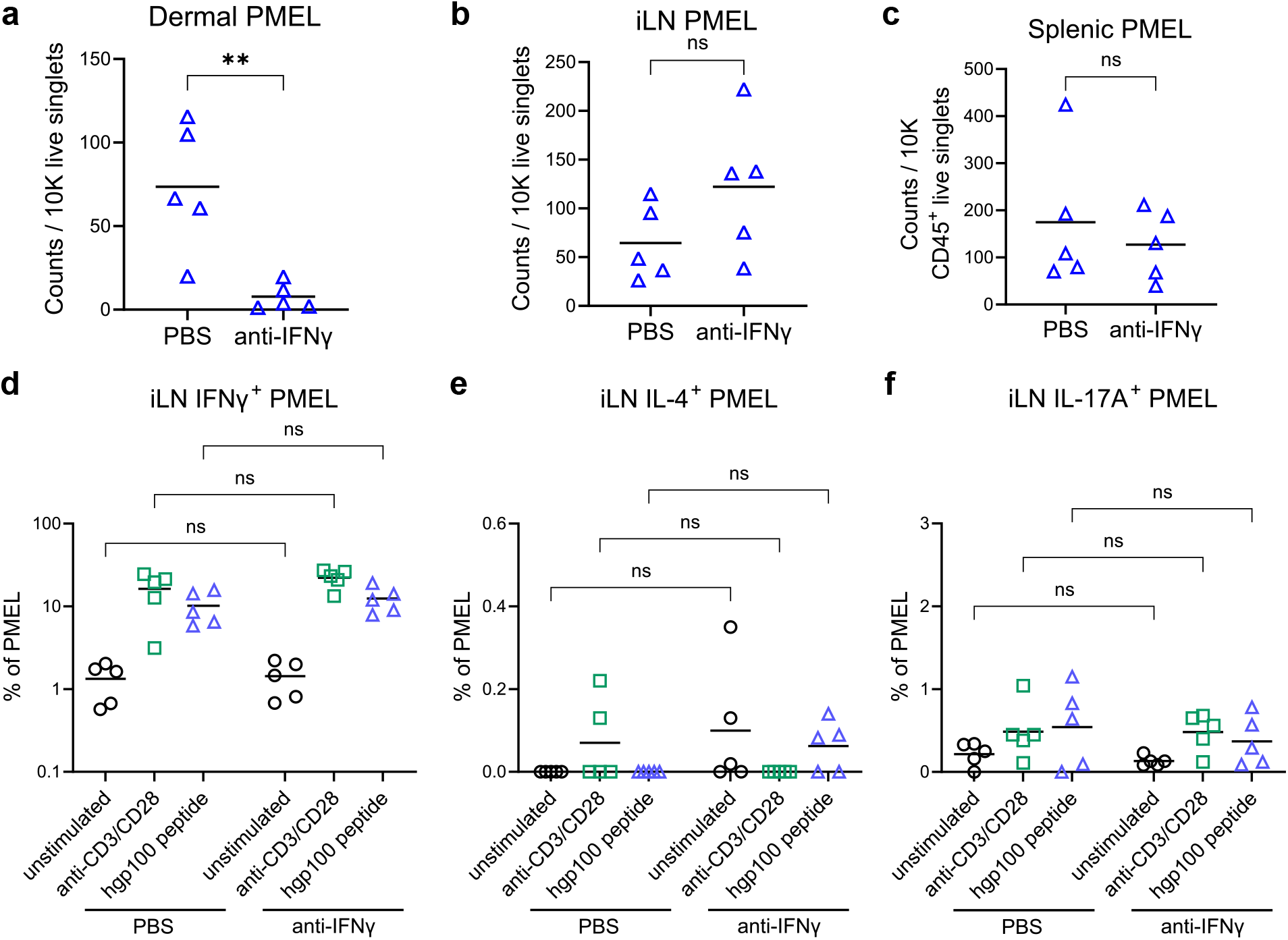
Systemic administration of anti-IFNγ does not affect PMEL T cell engraftment and cytokine production profile in lymphoid organs. Thy1.1^+^ PMEL T cell numbers in the (**a**) tail dermis (**b**) inguinal lymph nodes (iLN) and (**c**) spleen of vitiligo mice after 7 weeks of PBS or 250 ug anti-IFNγ weekly treatment. The iLN cell suspensions were restimulated overnight with anti-CD3 (10 ug/ml, plate-bound)/CD28 (2 ug/ml, soluble), or 10 uM human gp100_25-33_ peptide. PMEL production of (**d**) IFNγ, (**e**) IL-4, and (**f**) IL-17A were measured by intracellular staining and flow cytometry analysis. Data means are indicated from n=5 mice per group. Student’s *t*-test (**a-c**) or two-way ANOVA, Bonferroni post *hoc* test (**d-f**); ns, not significant, *\*\*P* < 0.01. All results were reproduced in another independent experiment.

**Fig. S2.**
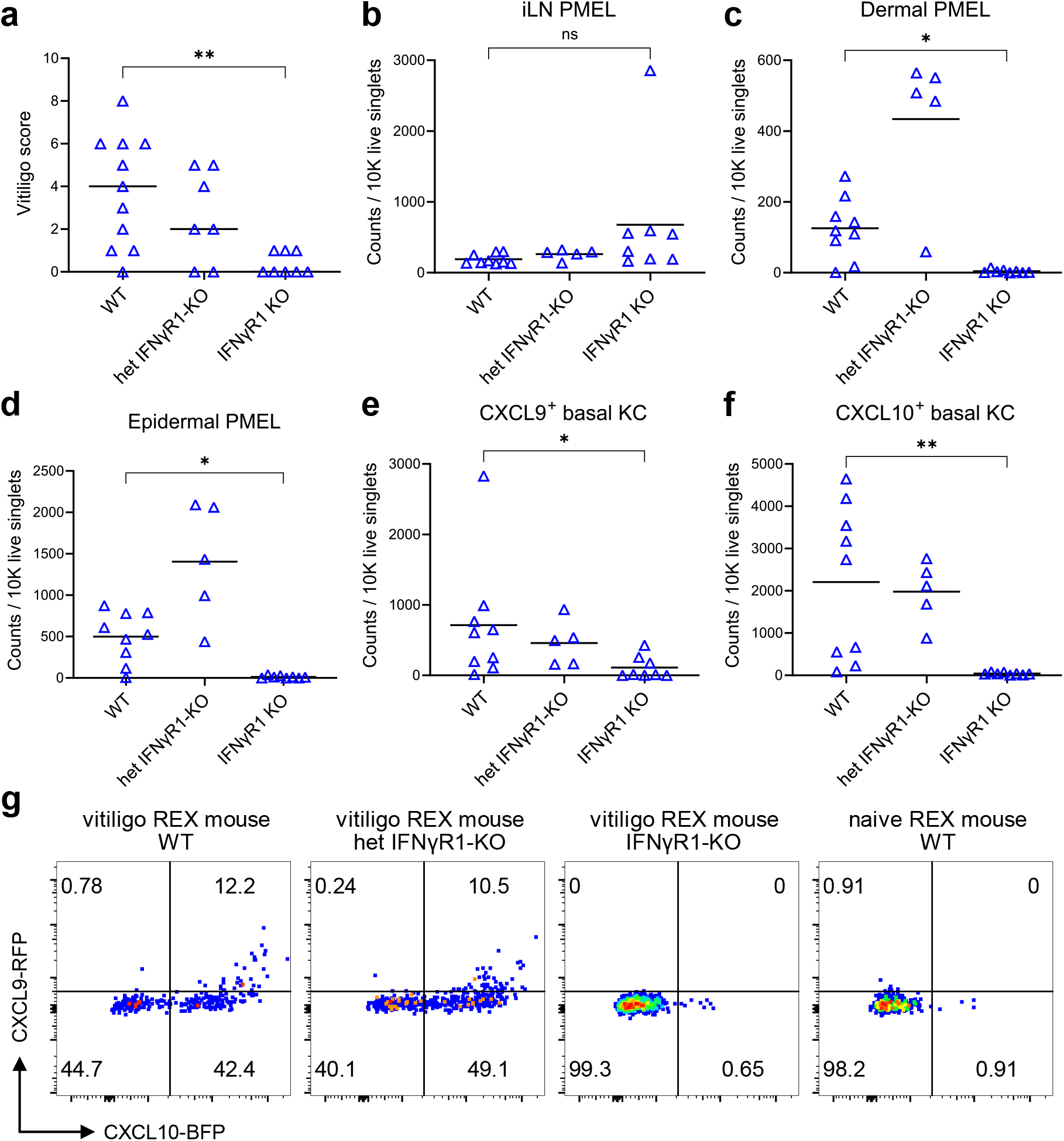
Host IFNγR1-deficiency abolishes basal keratinocyte CXCL9/CXCL10 production and skin PMEL T cells recruitment in the rVV-hPMEL vitiligo mouse model. Krt14-Kitl* REX WT or Krt14-Kitl* REX IFNγR1-KO mice received rVV and GFP^+^ PMEL T cells via adoptive transfer for vitiligo induction. (**a**) Vitiligo scores taken after 7 weeks, as well as quantification of GFP^+^ PMEL T cell numbers in the (**b**) inguinal lymph nodes, (**c**) tail dermis, (**d**) epidermis. (**e**) CXCL9 RFP and (**f**) CXCL10 BFP reporter expression of tail CD49f^+^ basal keratinocytes (KC) and (**g**) representative flow cytometry plots. Data means are indicated, except in **a** which are medians. One-way ANOVA, Bonferroni post *hoc* test; ns, not significant, \**P* < 0.05, \*\**P* < 0.01. Data were compiled from two independent experiments. WT, n=9 mice; het IFNγR1-KO, n=5 mice; IFNγR1 KO, n=8 mice.

**Fig. S3.**
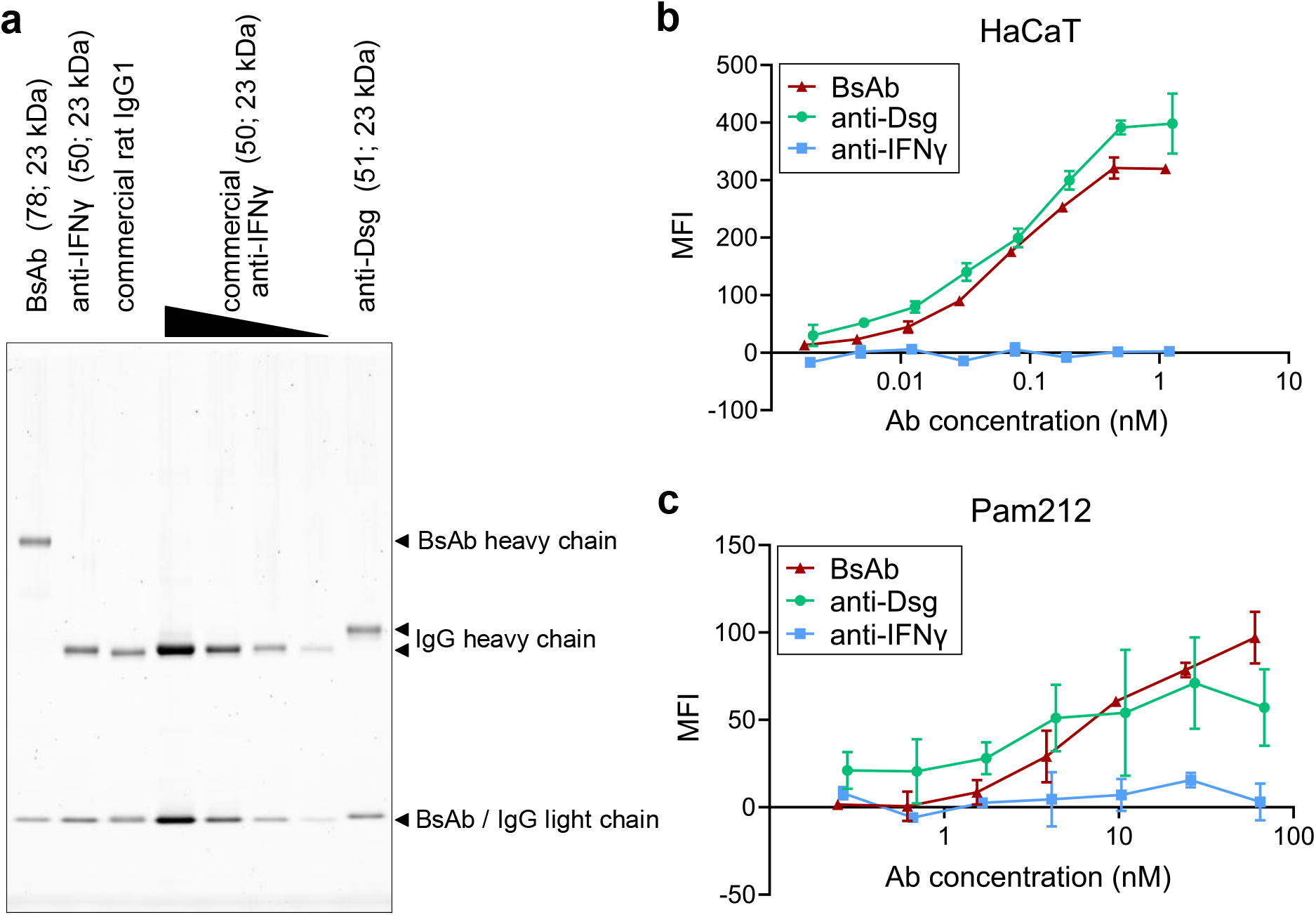
Dsg-IFNγ bispecific antibody binds human and mouse keratinocyte cell lines. (**a**) Purified antibodies were subjected to SDS-PAGE within reducing buffer and Krypton fluorescent protein stain. The theoretical molecular weights are indicated as (heavy chain; light chain). Antibody binding to (**b**) human HaCaT and (**c**) mouse Pam212 keratinocyte line. Ab, antibody. Data are expressed as mean±SD. All results were reproduced in another independent experiment.

**Table S1.** Antibody endotoxin level.

|  | Mean (EU/mg) | SD |
| --- | --- | --- |
| **Commercial anti-IFNγ** | 0.422 | 0.426 |
| **Purified anti-IFNγ** | 0.035 | 0.014 |
| **Purified BsAb** | < 0.002 | Not determined |

**Fig. S4.**
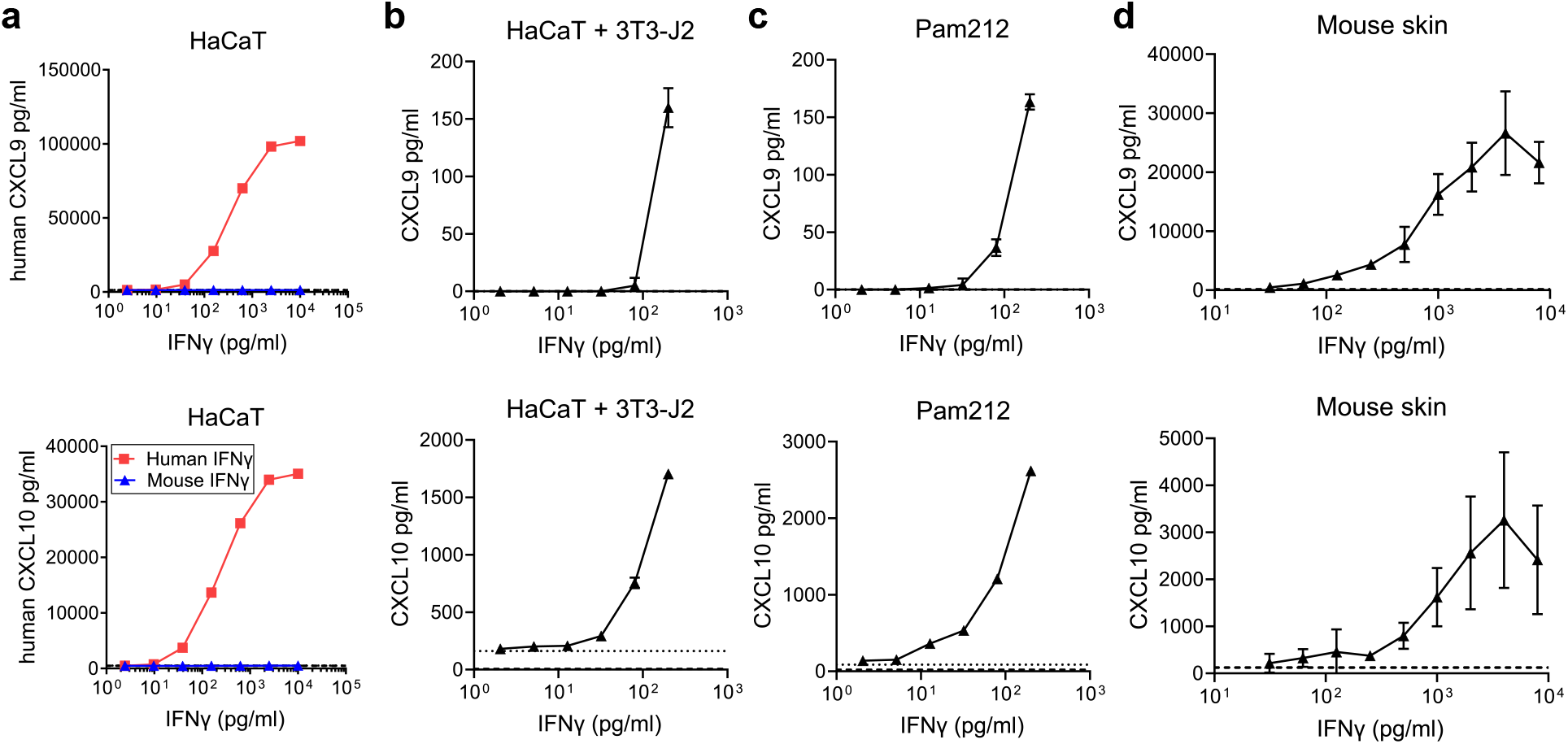
IFNγ dose-dependent chemokine production in the three *in vitro* and *ex vivo* skin models. (**a**) Confluent culture of HaCaT cells were stimulated with 10 ng/ml human TNFα and a varying concentration of human or mouse IFNγ for 18 hrs. The supernatants were collected and human CXCL9 and CXCL10 were quantified by ELISA. (**b**) HaCaT and 3T3-J2 co-culture, (**c**) Pam212 confluent culture, and (**d**) 4 mm tail skin punch biopsies from treatment-naïve mice were stimulated with 10 ng/ml mouse TNFα and a varying concentration of mouse IFNγ for 18 hrs. The supernatants were collected and mouse CXCL9 and CXCL10 were quantified by ELISA. Dashed and dotted lines indicate the chemokine level of medium alone and TNFα stimulation alone, respectively. Data are expressed as mean±SD. All results were reproduced in another independent experiment.

**Fig. S5.**
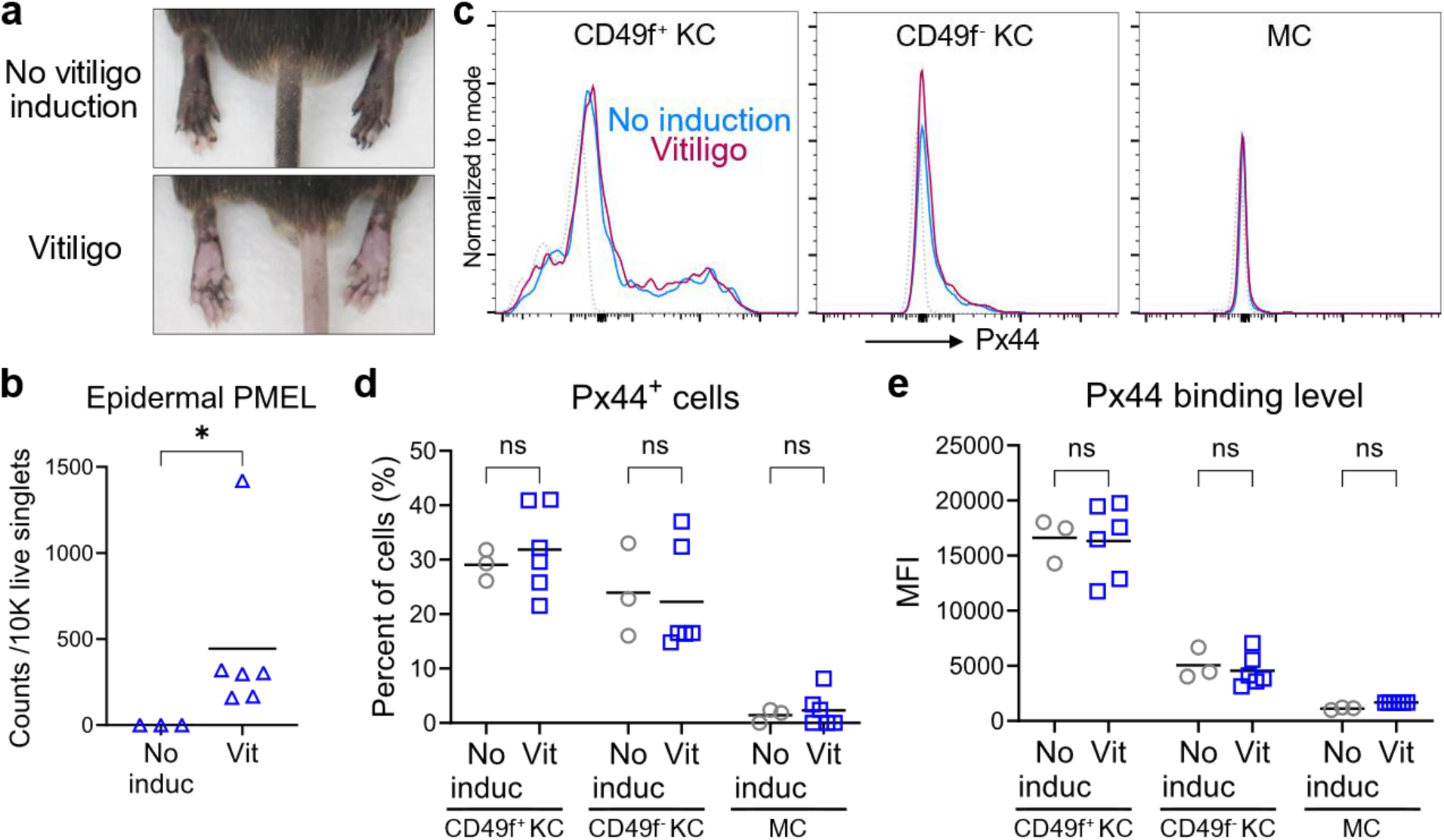
Anti-Dsg antibody Px44 binds healthy and vitiligo lesional skin at a similar level. (**a**) Representative image of the hind footpads harvested from 10-week-post-induction vitiligo mice or age-matched Krt14-Kitl* mice without vitiligo induction. Epidermal single-cell suspensions were prepared from resected tissue and analyzed by flow cytometry. (**b**) Quantification of epidermal Thy1.1^+^ PMEL T cells. (**c**) Px44 binding capacity was determined using biotinylated anti-Dsg or rat IgG1 followed by streptavidin-AF647 staining. Representative Px44 binding level histograms of CD49f^+^ basal and CD49f^-^ suprabasal keratinocytes (KC, gated by CD45^-^, keratin^+^), and melanocytes (MC, gated by CD45^-^, CD117^+^, CD49f^-^, keratin^-^). Isotype control staining is indicated by dotted lines. (**d**) Px44^+^ cell percentage. (**e**) Mean Px44 binding level. The MFI by rat IgG1 isotype staining was subtracted from each cell type. Data means are indicated from n=3 (no induction) or n=6 (vitiligo) mice per group. One-tailed Welch’s *t-* test (**b**) or two-way ANOVA, Bonferroni post *hoc* test (**d** and **e**); ns, not significant, \**P* < 0.05. All results were reproduced in another independent experiment.

**Fig. S6.**
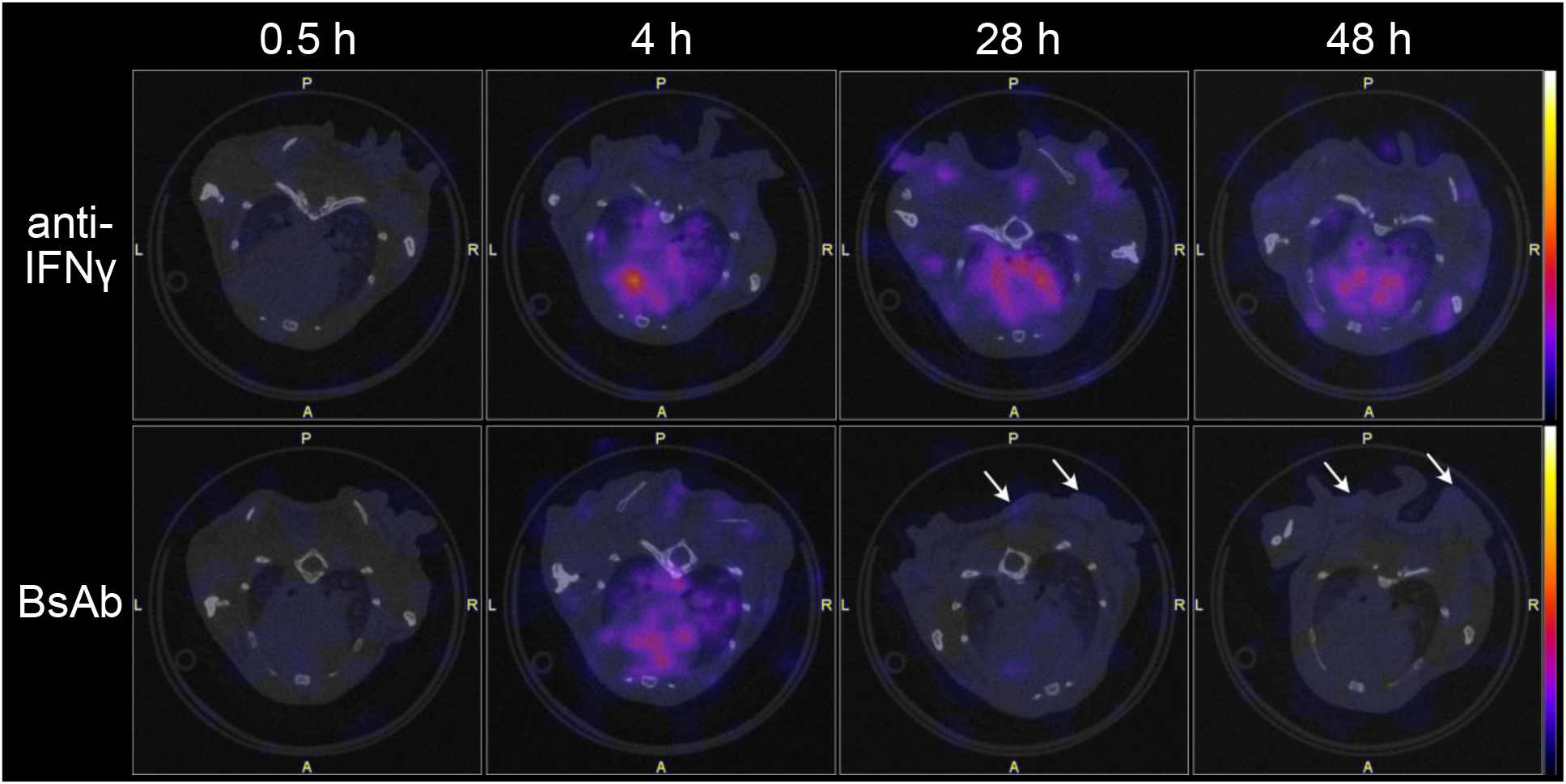
Time course SPECT/CT scan in transverse plane of the iodine-125 labeled antibody recipient mice. Krt14-Kitl* mice received 7.1 pmol ^125^I-labeled antibodies by right footpad injection. Images show the transverse plane of the chest. White arrows indicate ^125^I-BsAb signal from the skin at 28 hrs and 48 hrs post-injection.

**Table S2.** Degree of antibody iodine-125 labeling.

|  | Mean (Ci/mol) | SD |
| --- | --- | --- |
| **anti-IFNγ** | 5.10E+06 | 9.94E+05 |
| **BsAb** | 5.52E+06 | 7.81E+05 |

**Table S3.** Antibody half-lives* (hrs) in mouse tissues after single dose of footpad injection.

| Location | Antibody dose (pmol) | Antibody labeling type | Measurement methods | anti-IFNγ Mean (95% CI) | BsAb Mean (95% CI) |
| --- | --- | --- | --- | --- | --- |
| **Footpad** | 7.1 | Iodine-125 | SPECT/CT scan | **7.15**<br>(6.10, 8.14) | **15.62**<br>(12.23, 19.78) |
| **Disseminated body (excl. footpad)** | 7.1 | Iodine-125 | SPECT/CT scan, radioisotope calibrator | **88.82**<br>(68.46, 159.00) | **13.30**<br>(6.68, 30.47) |

\*Half-lives were derived from fitting the data to first order elimination kinetics

**Fig. S7.**
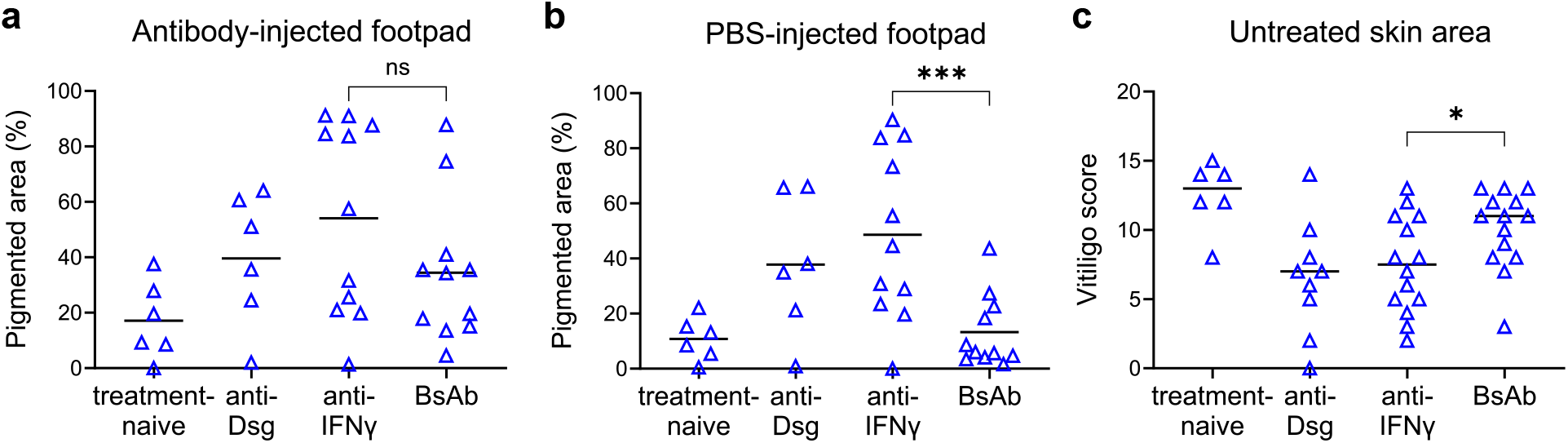
The BsAb does not have a systemic therapeutic effect at the untreated skin area. After receiving PMEL T cells and BMDC co-transfer to induce vitiligo, mice were treated with antibody and PBS to the right and left footpad, respectively, every other day for 5 weeks. Pigmented skin area was quantified by ImageJ in the (**a**) antibody-injected footpad and (**b**) PBS-injected footpad. (**c**) Total vitiligo scores of the nose, ears, tail, and PBS injected footpad combined. Data means are indicated, except in **c** which are medians. One-way ANOVA, Bonferroni post *hoc* test; ns, not significant, \**P* < 0.05, \*\*\**P* < 0.001. Data were compiled from two independent experiments.

**Fig. S8.**
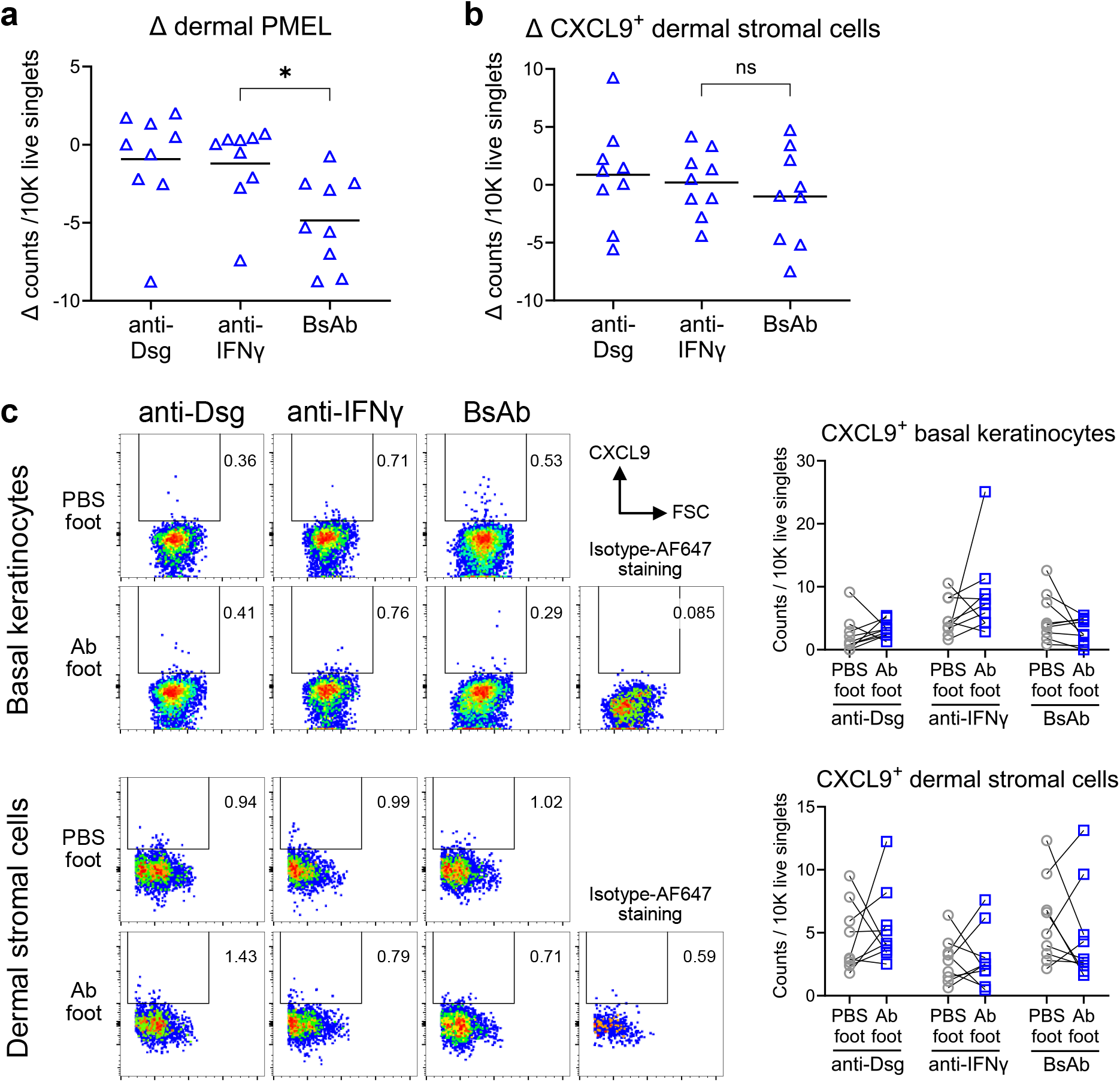
The effect of the BsAb in the dermis of treated footpad. Krt14-Kitl* mice received antibody treatments to the right hind footpad and PBS to the left footpad every other day for 5 weeks following vitiligo induction. (**a**) Δ footpad dermal Thy1.1^+^ PMEL T cell number. (**b**) Δ footpad CXCL9^+^ dermal stromal cells (gated by CD45^-^, CD117^-^, keratin^-^) number. (**c**) Representative CXCL9 intracellular stainings of the footpad basal keratinocytes (gated by CD45^-^, CD117^-^, CD49f^+^ keratin^+^) from epidermis samples and the dermal stromal cells from dermis samples. Data means are indicated from n=9 mice per group. One-way ANOVA, Bonferroni post *hoc* test; ns, not significant, \**P* < 0.05. All results were reproduced in another independent experiment.

**Fig. S9.**
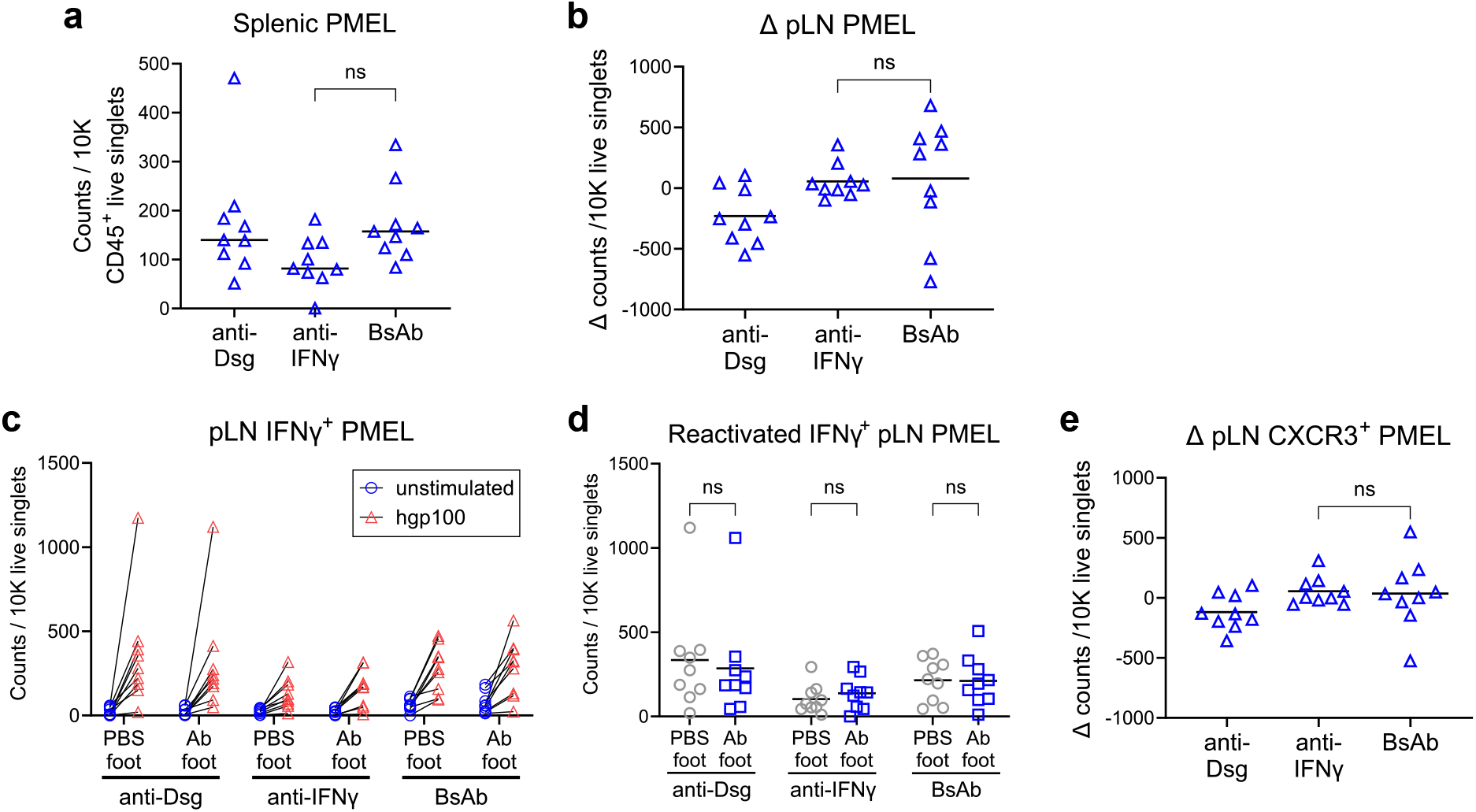
Unilateral popliteal lymph node antibody drainage does not account for the asymmetrical footpad skin inflammation status. Krt14-Kitl* mice received footpad injection treatment every other day for 5 weeks following vitiligo induction. (**a**) Splenic Thy1.1^+^ PMEL T cell engraftment. (**b**) Δ PMEL T cell number in the popliteal lymph nodes. The popliteal single-cell suspensions were further restimulated overnight with human gp100_25-33_ peptide and measured for (**c**) IFNγ^+^ PMEL T cell number before and after the restimulation. (**d**) The mathematical difference in the number of IFNγ^+^ PMEL T cells between pre- and post-peptide restimulation. (**e**) The CXCR3^+^ PMEL T cell number post-peptide restimulation. Data means are indicated from n=9 mice per group. Two-way ANOVA with Bonferroni post *hoc* test (**d**) or one-way ANOVA with Bonferroni post *hoc* test; ns, not significant. All results were reproduced in another independent experiment.

## Supplementary Materials

### List of supplementary materials

#### Supplementary videos

Video S1. anti-IFNg_0.5h

Video S2. BsAb_0.5h

Video S3. anti-IFNg_4h

Video S4. BsAb_4h

Video S5. anti-IFNg_28h

Video S6. BsAb_28h

Video S7. anti-IFNg_48h

Video S8. BsAb_48h

